# Meta-analysis of macrophage nanoparticle targeting across blood and solid tumors using an eLDA Topic modeling Machine Learning approach

**DOI:** 10.1101/2023.06.29.547096

**Authors:** Chloe Brown, Colette Bilynsky, Melanie Gainey, Sarah Young, John Kitchin, Elizabeth Wayne

## Abstract

The role of macrophages in regulating the tumor microenvironment has spurned the exponential generation of nanoparticle targeting technologies. With the large amount of literature and the speed at which it is generated it is difficult to remain current with the most up-to-date literature. In this study we performed a topic modeling analysis of the most common usages of nanoparticle targeting of macrophages in solid tumors. The data spans 20 years of literature, providing an extensive meta-analysis of the nanoparticle strategies. Our topic model found 6 distinct topics: Immune and TAMs, Nanoparticles, Imaging, Gene Delivery and Exosomes, Vaccines, and Multi-modal Therapies. We also found distinct nanoparticle usage, tumor types, and therapeutic trends across these topics. Moreover, we established that the topic model could be used to assign new papers into the existing topics, thereby creating a Living Review. This type of meta-analysis provides a useful assessment tool for aggregating data about a large field.

## Introduction

Macrophage nanoparticle targeting in cancer immunotherapy is an exploding field. Tumor associated macrophage (TAMs) are a prime target for biologists who seek to understand their role in tumor progression and immune evasion. Likewise, they have been a target for engineers who design therapeutics to reprogram TAM functions. The enormous amount of literature generated spans multiple scientific disciplines, institutions and geographical regions. In addition, literature is accessed across numerous databases. Moreover, differences in disciplines can lead to different nomenclatures which may limit the accuracy of traditional literature search strategies. All of these complexities can make it difficult for any single research to stay current.

A scoping review is a powerful tool that gives researchers a broad picture of a field by extracting data from literature related to a specific research question. To accomplish this goal, papers are processed through multiple phases of searching, screening, and data extraction. By reviewing a large volume of work, researchers can then review an entire volume of literature and obtain a qualitative and quantitative view of a field. This systematic approach has other advantages as well. It can reduce bias that is introduced when reviewing a smaller number of papers. Furthermore, by utilizing a protocol for screening papers, it can help identify emerging research trends.

One of the challenges of a scoping review is the time and work needed to screen and extract data from thousands of papers. Machine learning, and specifically natural language processing, have been used to aid in multiple phases of a scoping review[1]–[3]. Previously, topic modeling, a form of unsupervised machine learning, was used to create an overview of the types of papers in a literature review based on their abstracts [4]. These include but are not limited to cancer immunotherapy [5], emergency medicine [6], and HIV-AIDS research [7].

In this paper, we generate a scoping review about macrophage targeting cancer therapy using the Gensim Ensemble Latent Dirichlet Allocation (eLDA) topic modeling algorithm [8], [9]. The articles used in this project were the result of a full-text screening process using pre-defined eligibility criteria, previously described in “Scoping Review of Pre-clinical and Translational Studies on Macrophage Polarization in Nanoparticle–based Cancer Immunotherapy” [10]. Additionally, using this model, we demonstrate how LDA topic models can be used to create a “living scoping review” where new articles can be assigned to existing topics.

## Materials and Methods

### Datasets and Code

The datasets used to create this model can be found in the supplemental information. Additionally, a Jupyter notebook containing sample code to create the model and figures can be found in the supplemental material. Finally, the files that contain the model can be found in the supplemental material.

### Obtaining the Dataset

The dataset for this review was obtained using guidance from several scoping review methodology resources [8,9]. Scoping reviews are used to obtain a near-comprehensive overview of a particular body of literature with a well-defined scope. A systematic approach to searching for and identifying relevant articles is used, with the aim of minimizing selection bias. With this approach in mind, a protocol was developed and registered on the Open Science Framework [10]. A search string was crafted by two of the authors who are information specialists with experience in evidence synthesis methods (MG, SY) to search for articles containing information about nanoparticles and macrophage polarization in a cancer context **(Supplemental File 1)**. We searched the following bibliographic databases: Web of Science Core Collection (including Science Citation Index-Expanded, Social Science Citation Index and Emerging Sources Citation Index), Scopus, IEEE Xplore Digital Library, Medline (PubMed), and Biotechnology & BioEngineering Abstracts (ProQuest). Additional articles were found by hand-searching journals, references from related literature reviews, and Google Scholar. The database searches were run between April 23, 2020 and October 20, 2020. We chose to only include articles published in 2000 or later due to the large growth of research in cancer nanotheranostics in the last twenty years. Due to the large number of included articles and limited resources, we did not conduct forward or backward citation searching.

The articles, hereafter referred to as records, underwent the first few stages of a scoping review, including de-duplication, title and abstract screening, and full text screening [11]. In the de-duplication phase, records that appeared multiple times in the dataset were consolidated into one entry. This work was done in Zotero.

In the title-abstract screening phase, records were labeled as “include” or “exclude” based on a set of pre-defined criteria [10]. These criteria included whether the study was carried out in a cancer context, if the study included information about the characterization and use of nanoparticles, and whether a study may contain information about macrophage polarization. To be fully included or excluded, two independent reviewers needed to have identical labels. In the event of a conflict, a third expert reviewer determined the final label.

During the full text screening, records were again labeled “include” and “exclude” based on more refined criteria. As with the previous screening stage, inclusion decisions needed to be consistent between two independent reviewers, with conflicts being resolved by a third expert reviewer. Papers that were excluded in this section were also labeled with an exclusion reason. The title-abstract and full text screening phases were carried out in Sysrev [12].

We used Sysrev’s built-in machine learning capabilities to automatically exclude articles in our large dataset [12]. We first trained the Sysrev inclusion prediction algorithm by manually screening half (7,482) of the records in the initial title-abstract phase of screening. We then used its prediction to automatically exclude records with a prediction value of less than 40% (3,460 records). We found that 14 of the 7,482 articles (0.19%) in the training dataset had been incorrectly classified for exclusion by the prediction algorithm when using a prediction value of less than 40%. These records were manually included in the full-text screening phase of the project. Records identified from Google Scholar, Biotechnology & BioEngineering Abstracts (ProQuest), and hand searching were not used to train the algorithm and were manually screened for inclusion at a later stage of the project.

The topic model used in this study was created using 854 of the abstracts of papers included after the full text screening **(Supplemental File 2)**. An additional 95 abstracts retrieved using the same search engine strategies to create a living scoping review.

### Data Pre-Processing

The dataset went through multiple phases of pre-processing including converting all words to lowercase, tokenization, removing stop words, and stemming.

#### Tokenization and Converting to Lowercase

The NLTK RegEx tokenizer was used to tokenize the document set [3]. We used the regular expression r’\w+’ which matches all Unicode word characters. The tokens for each document were then converted to lowercase.

#### Stop Words

A list of stop words was created in multiple steps to remove from the data set. First, a list of common English stop words was imported from NLTK [3]. Next, a list of stop words was created based on the search string. These included words like “cancer”, “nanoparticle”, and “macrophage” which were common in the dataset, even though they are not common in the English language. Finally, words that occurred in one abstract or 90% of abstracts were removed. Sometimes, a domain-specific list of stop words is needed, such as when an abstract set has a narrow scope [4].

#### Stemming

The abstracts were stemmed using the NLTK Porter Stemmer [3]. We chose to stem the abstracts rather than lemmatize them because many words in the dataset are very uncommon in colloquial English. Therefore, lemmatization would not work as well on this dataset.

### LDA Algorithms

The traditional LDA model generated noisy, incoherent topics and this result was insensitive to hyperparameter tuning. Therefore, we attempted to use a modified topic modeling algorithm that was developed to decrease the noise associated with the random allocation of topics when initializing LDA [13]. Gensim Ensemble Latent Dirichlet Allocation (eLDA) uses the DBSCAN algorithm on a collection of LDA topics to distinguish between stable and noisy topics [14]. We found that eLDA had superior performance to the traditional LDA model and used this technique for the scoping review [15]. In addition, we have made the code available for public use **(Supplemental File 3)**.

The Gensim eLDA module requires the parameters corpus, which is a bag of words model of the documents set, and id2word, a mapping of all words to an integer identity. The dictionary was created using the Dictionary module from the Gensim corpora module. The bag of words model of the document set, the corpus, was created by applying the doc2bow function from the dictionary to every abstract in the cleaned dataset.

### Evaluating the Topic Model

The topic model was evaluated both quantitatively and qualitatively. The model was evaluated quantitatively using the metric average topic coherence, which was measured using the Gensim Coherence Model module [15]. The topics created by the models were also evaluated qualitatively for coherence with individuals with domain knowledge in the field. This was done by evaluating the top 30 highest weighted words from each topic distribution as defined by the LDA model. Finally, we looked at the 100 most frequent terms to look for trends within each topic based on previous domain knowledge.

### LDA for a Living Review

We used the 95 new abstracts to demonstrate a “proof-of-concept” for a living review. To do this, we first pre-processed the abstracts as described above and created a “bag-of-words” model of each document. We then used the eLDA model to extract topics for each model. LDA, and by extension eLDA, has the ability to extract topics from documents that were not previously seen in the dataset, a strategy that has previously been used in classification studies [8], [16], [17]. In this instance, we did not update the model to incorporate the new documents.

## Results and Discussion

### Overview of the Scoping Review and the generated dataset

The searches of bibliographic databases, handsearching and Google Scholar resulted in 28,217 records. After deduplication and removing retracted records, 15,374 were screened with 859 records included in the final dataset (Figure 1). 854 records were used to train the topic model and 95 new records were used to generate a living scoping review. Note that the 95 new records did not undergo the screening process.

**Figure 1:**
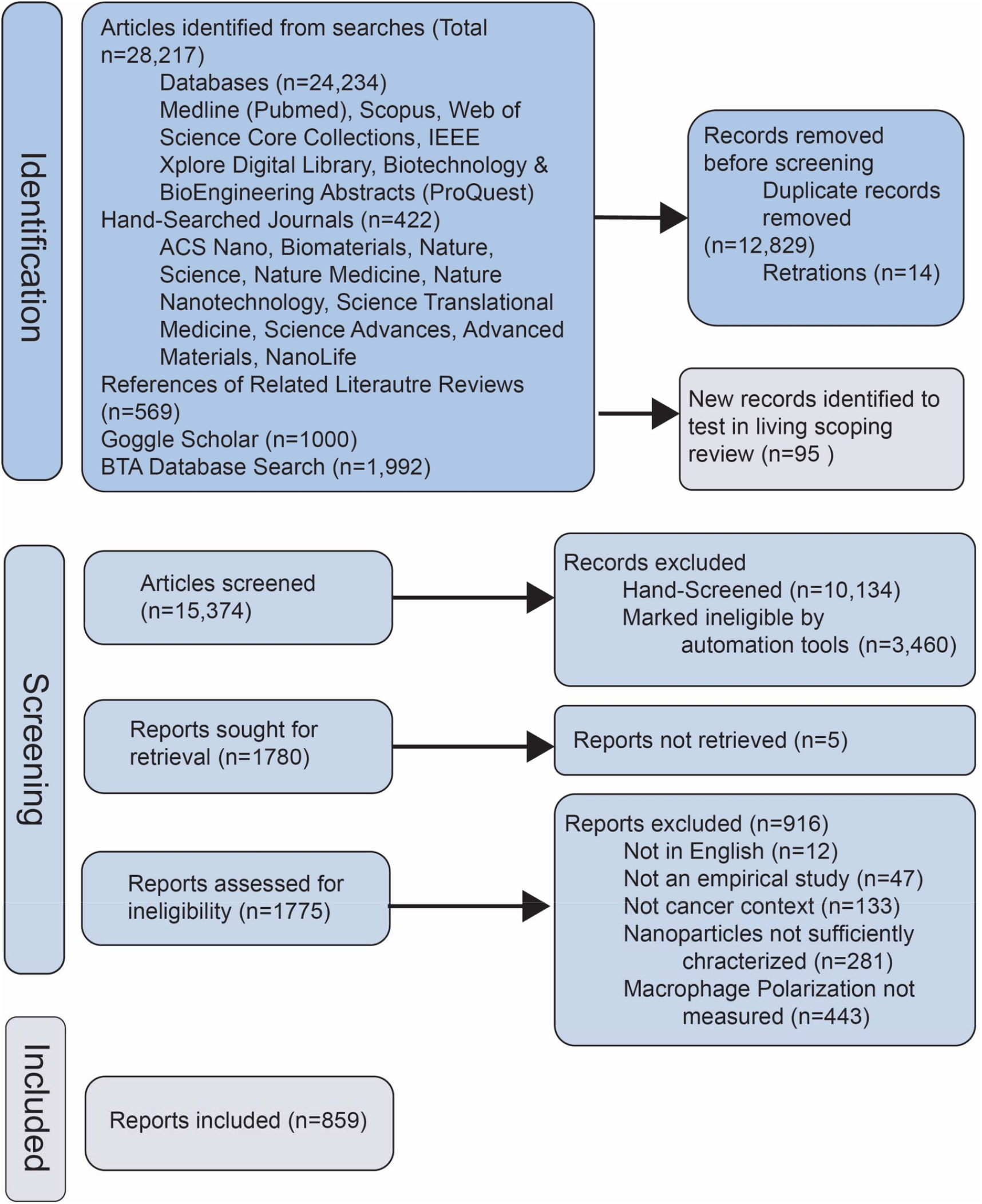
Obtaining the Dataset Diagram of the scoping review methodology. Flow chart outlining the number of records from article identification, screening, and inclusion into the final data set used in the eLDA topic model. Of the 28,217 articles identified, 859 articles that met the inclusion criteria. 95 newly identified records were used in the Living Review model. Adapted from [18].

Before creating the topic model, analyses were carried out to characterize the dataset, which can be found in **(Figure 2)**. First, we investigated the trends over time in the publications **(Figure 2A)**. We observed two inflection points of growth in the field of macrophage cancer immunotherapy. These represent two growth fields in the immunotherapy field. In 2011 Ipilimumab, the immune checkpoint inhibitor, was approved by the FDA for use in melanoma [19]. Moreover, the 10 year sequel to the original Hallmarks of Cancer included the role of the immune system as a principal biological component in cancer progression and treatment [20]. In 2017, the first CAR T therapy was approved by the FDA [21]. During this time period, it became increasingly apparent that the immune system played a crucial role in finding effective treatments for cancer. Both of these advances utilize T-cells, which is why they have been most useful for tumors with high T-cell invasion as well as liquid tumors [19]. Macrophages make up a large portion of the volume within tumors and contribute to the immunosuppressive environment that prevents T-cell infiltration [22]. Perhaps for this reason, macrophages specifically became a target for cancer nanotherapeutic research, resulting in exponential growth in this field.

**Figure 2:**
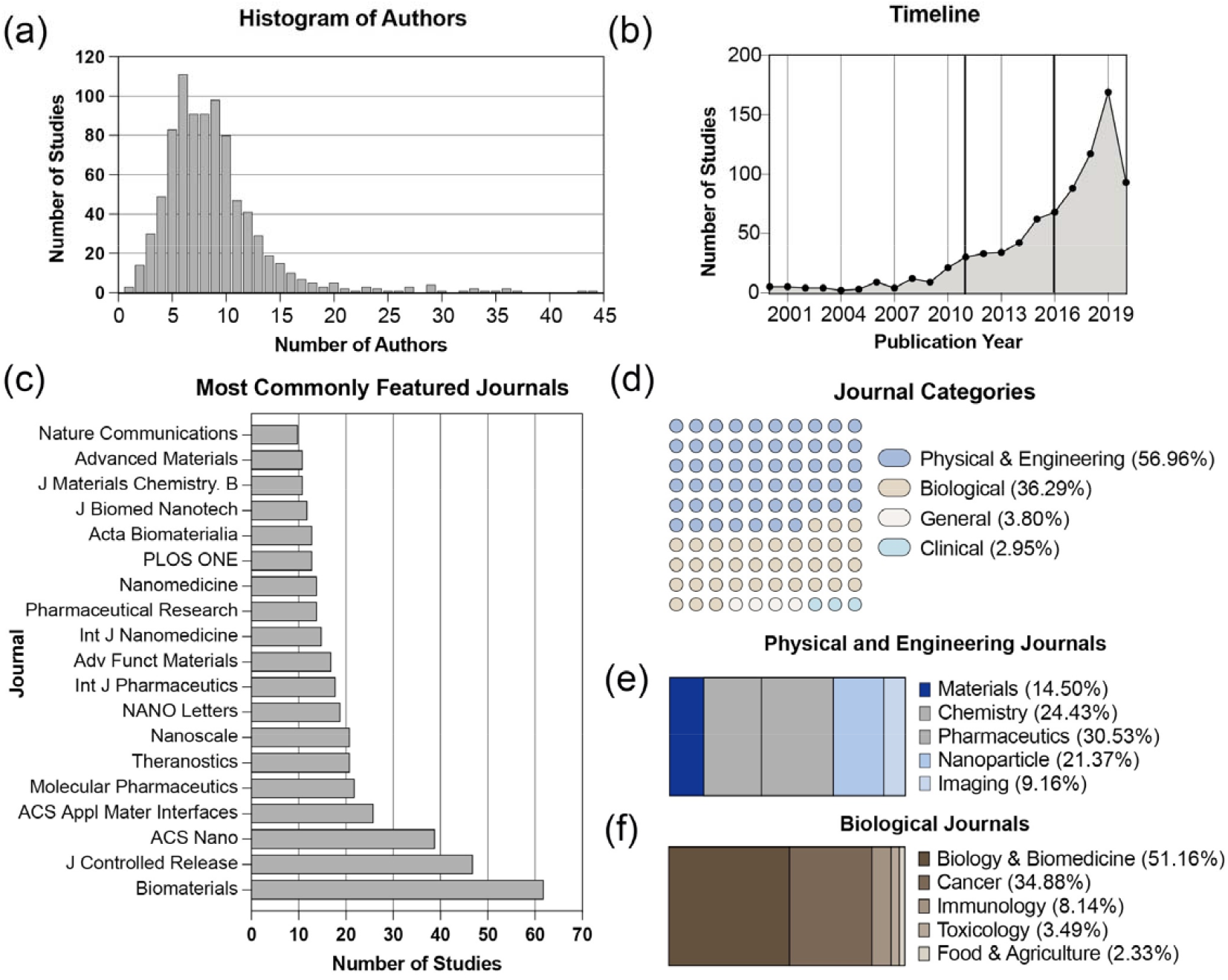
Characterizing the Dataset: (a) Distribution of number of authors in the dataset. (b) Number of documents in the dataset over time. Gray lines indicate important advancements in the field. (c) Most common journals in the dataset. (d) Distribution of journal topics included in the dataset. (e) Breakdown of the journal scopes within (e) physical and engineering journals and (f) biological journals.

Further bibliographic analysis demonstrated other publishing trends within the field. The median number of authors was about 8, suggesting that this type of research frequently requires a larger collaborative team, since immunomodulatory nanotherapeutics will often need a multidisciplinary approach (Figure 2B). We did not extract data related to author gender and institution because the sociological expertise needed to properly extract and analyze these features was viewed to be out of scope. Overall, the articles were published in an array of journals with Biomaterials, Journal of Controlled Release, and ACS Nano being the most represented (Figure 2C). All three of these journals emphasize the multidisciplinary nature of their publication: physics, chemistry, and biology.

### Characterization of the Topic Model

The topic model was evaluated both qualitatively and quantitatively to evaluate the accuracy of the model. Average topic coherence was used as a quantitative evaluation [23]. The LDA hyperparameters and number of topics are typically tuned using the coherence score. However, we found that varying these hyperparameters did not have a significant impact on the coherence score. Instead, we chose hyperparameters to create a model that fit our purposes. The Cv coherence score for this model was 0.589. A visualization of the topic model can be found in **(Figure 3)**, which was created using pyLDAvis [24].

**Figure 3:**
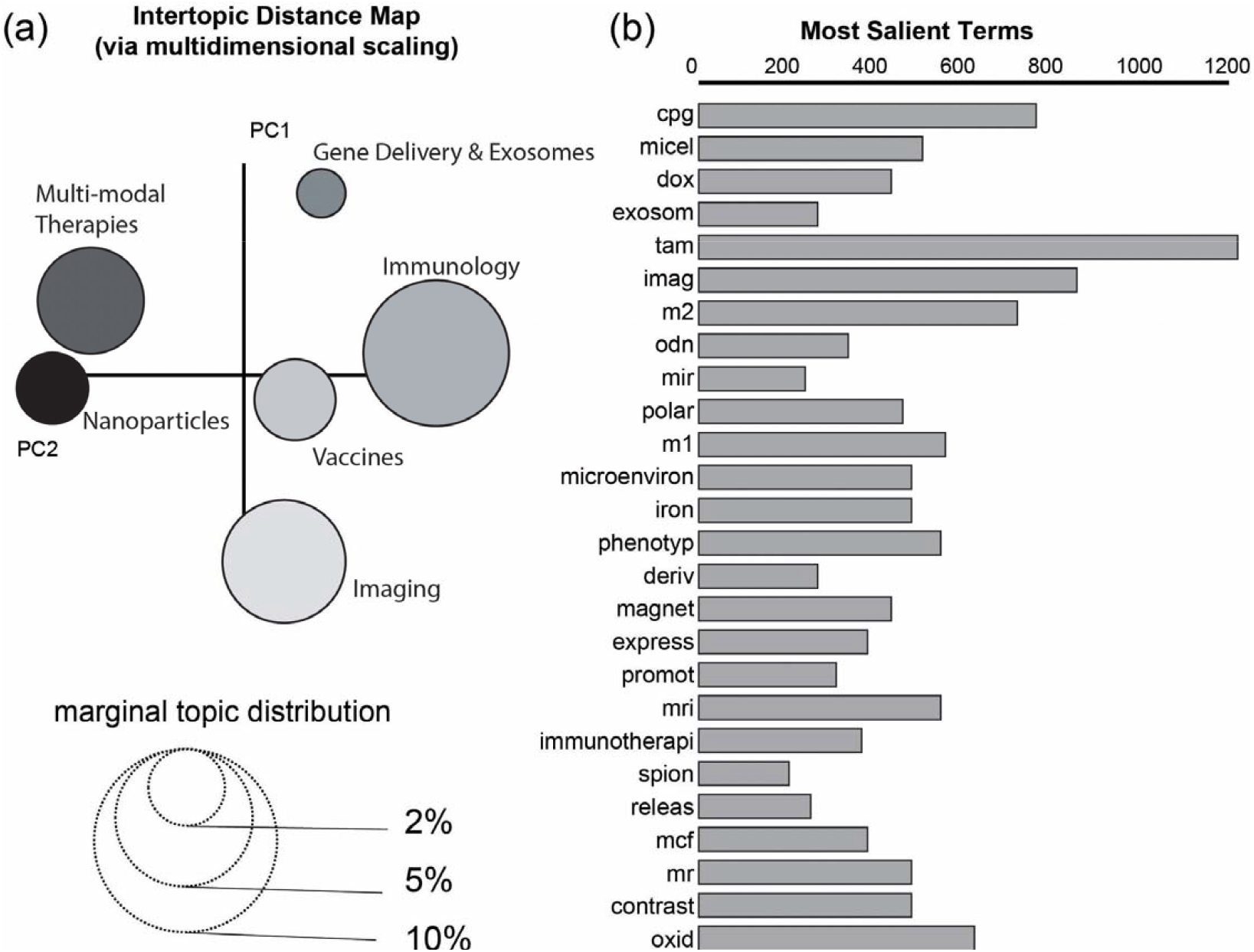
Creating the Ensemble LDA Topic Model. (A) Intertopic Distance Map of LDA topics. Size of the circles represents the relative number of articles within each topic while the distance between circles represents similarity between topics. Marginal topic distribution for each topic indicates the percentage of the dataset that belong to each topic. (B) Most Salient Terms. These are the thirty most frequent terms used to create the topic model. This figure was generated using the pyLDAvis library for topic model visualization in Python [24].

For the qualitative evaluation, the topic model was determined to be successful if it produced topics that made human sense. That is, a successful topic model had topics with distinct themes based on the top words and documents assigned to them. The topics were named in accordance with these themes. The topics were qualitatively assigned names based on abstract contents as follows: Immune and TAMs, Nanoparticles, Imaging, Gene Delivery and Exosomes, Vaccines, and Multi-modal Therapies.

Although the eLDA algorithm was effective in reducing the generation of incoherent topics, it did not result in reproducible topics over multiple runs [23], [24]. This is because the LDA algorithm which is the basis for eLDA uses a random allocation to initially assign topics. The random allocation in the model can return different topics when given the same parameters. When utilizing this strategy, we compiled the model multiple times to determine the most persistent topics to represent the dataset. We calculated the average topic coherence for each topic model created and found that it did not vary significantly between runs.

Qualitatively, we found the model most frequently returned to Immunology and Nanoparticle Topics. The remaining topics appeared throughout the model simulations but not as consistently. Imaging and Multi-model Therapies were usually found in the Nanoparticle Topic while, Vaccine, Gene Delivery and Exosomes topics were contained within the Immunology Topic. This seemed reasonable given the topic proximity on the Intertopic Distance Map (Figure 3A). When evaluating the topic models qualitatively, the models produced using eLDA still make more “human sense” than the topic models created with LDA alone with this dataset. Therefore, while the eLDA algorithm did not create entirely reproducible models, it did reliably create meaningful topics. The eLDA model therefore discussed throughout the rest of this manuscript is the one that we found to best characterize the major and minor features of the dataset.

The Immune and TAMs section had the largest distribution of documents, this was expected because the searching and screening phases of the scoping review targeted documents that discussed macrophage polarization in the context of cancer immunotherapy. Interestingly, the smallest topics were Nucleic Acids and Exosomes and Nanoparticles.

The most salient terms, the most frequent terms in the corpus, are shown in (Figure 3B). The bar length represents the term frequency in the entire dataset. The high prevalence of TAM is unsurprising, as the scoping review targeted studies in which nanoparticles interacted with tumor-associated macrophages (TAMs).

### Identification of distinct categories within the dataset

To evaluate the topic contents, we categorized the most frequent words that occurred within each topic. Each article was assigned into a topic if the weight for that topic was at least 0.40. These words were then categorized by theme and sorted by frequency within the dataset. **(Figure 4A)** summarizes the results of this analysis. In total, six categories of words were identified: immunology, nanoparticles, imaging, gene delivery and exosomes, vaccines, and multi-modal therapies.

**Figure 4:**
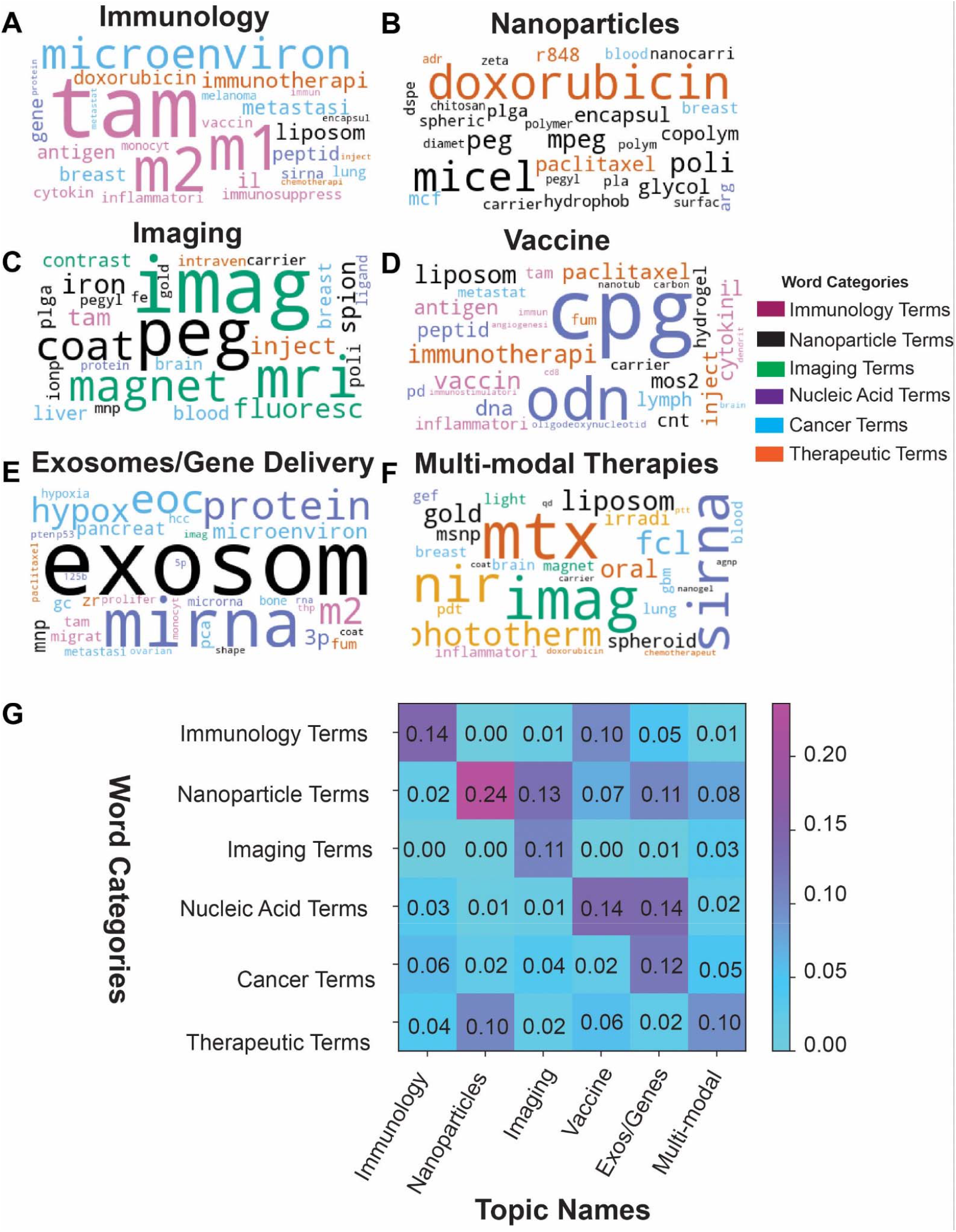
Dataset by word categories and topic names. (a) Word clouds by topic. Word size was determined by the frequency within the dataset while the colors were determined by their category. (b) Heat map of word categories within each topic.

Some topics were largely dominated by one type of word. The Immunology topic contained terms such as TAM, M1, and M2 listed prominently. Nanoparticle topics, which also had the highest weighted score of all the topics, predominantly contained words associated with nanoparticle formulation and design. However, the other topics were identified by the unique combination of two or more categories. The Imaging topic by contrast, was highlighted by “imaging” and “nanoparticle” terms. The Nucleic Acids and Exosome group was highlighted by “nanoparticle”, “nucleic acid” and “cancer” terms. Furthermore, these trends were quantitatively validated by the weight scores **(Figure 4B)**. The weight scores were calculated by dividing the frequency of words belonging to a given category by the total frequency of words in that topic. For example, in documents that were assigned to the nanoparticle topic, among all appearances of the 100 most frequent words, nanoparticle terms occurred approximately 24% of the time. It is important to note that words that were too vague to elucidate their meaning in the context of the documents were not included in this analysis, even if they did appear in the 100 most frequent terms. These include words like “activ”, “method”, and “cancer”.

It should also be noted that while more than one topic contained nanoparticles as a predominant word category, the kinds of nanoparticle terms that appeared in each topic were unique. For instance, the nanoparticle terms in the imaging topic such as “iron” and “gold” did not appear in the Nanoparticle topic (i.e., PLGA, micelle, chitosan) (Figure 4A).The nucleic acid word category with respect to the Vaccine topic contained “cpg”, “odn”, and “antigen” in comparison to the Nucleic Acids and Exosome group which contained “mRNA”, “microRNA”, and “3p”.

We also noticed that organs were more prominent based on the topic. For example, breast, liver, brain, and blood were key terms that appeared in the imaging topic. In nucleic acids and exosomes, pancreas, bone, prostate, and hepatocellular organs were prevalent.

#### Immunology

Papers in the Immunology topic tend to prioritize repolarization of tumor associated macrophages (TAMs). These papers describe the TAM phenotype and strategies to convert them into anti-tumoral pro-inflammatory macrophages. These strategies include cytokine treatment, microRNA gene delivery, and small molecule stimulation of pro-inflammatory transcriptional pathways such as NF-kB and TLR ⅞. Notably, these therapeutic cargoes are distinct from the types of therapeutics seen in the Nanoparticle Topic. Rather than attempting to induce death in cancer cells, these strategies are attempting to target TAM function, migration [25], [26], transcriptional activity, and antigen presentation [27]. This was a particularly strong topic within the data set, and consistently appeared during model reruns.

#### Nanoparticles

The nanoparticle topic showcases papers with a strong focus on nanoparticle characterization and development for anti-tumor therapy. While nanoparticles show up in other topics, papers that describe *in vivo* pharmacokinetic/pharmacodynamic (PK/PD), *in vitro* and *in vivo* cytotoxicity dose measurements, as well as structural and loading characterization are most likely to be found in this topic. Polymeric nanoparticles are the most common composition found followed by lipid/liposomes. In addition, chemotherapeutic drugs such as Paclitaxel, Docetaxel, and Doxorubicin feature prominently in this section. Such papers intend to overcome multidrug resistance [28]–[30]. This was another strong topic, persistently appearing alongside the immunology topic when the model was rerun.

#### Imaging

The imaging topic contains papers that use nanoparticles to visualize macrophages within tumors. The most prominent nanoparticles are metallic nanoparticles that provide great contrast in MRI imaging. Given the high prevalence of macrophages within solid tumors, these nanoparticles are also used to detect tumor metastasis [31]–[33]. Nanoparticles are conjugated with mannose [34]–[36] to enhance uptake in monocyte, macrophage, and dendritic cell populations. Moreover, these nanoparticles are used as theranostics (diagnostics + therapeutics). Many papers combine metal particles with therapeutics like curcumin [37] and a COX-2 inhibitor [38]. This topic has some overlap to the Multimodal Therapy Topic.

#### Gene Delivery and Exosomes

The Gene Delivery and Exosomes Topic featured papers that used extracellular vesicles (EV) drug delivery as well as intercellular signaling within tumors. Extracellular vesicles, which include exosomes, are released by cells, and can carry a wide array of cell constituents like DNA, RNA, and proteins [39]. EVs can be used as therapeutics, either as drug delivery vehicles or therapeutics as themselves. In these papers, most studies were using EVs to deliver nucleic acids, such as microRNA. The topic broadly focused on the delivery of biologics, besides microRNA, proteins like an anti-CD137 antibody and SIRP□ were also delivered using nanoparticles [40], [41]. Besides extracellular vesicles, this category also included nanoparticles with different surface functionalization aimed at improving cell-specific targeting: macrophage membrane, cancer cell membrane, and CD47 [42]–[44].

#### Vaccines

As the title describes, the topic contains vaccine strategies for macrophage reprogramming. Here, cargo strategies were not primarily chemotherapeutics but antigens and adjuvants designed to initiate pro-inflammatory activation. Lymph nodes were featured in this topic because of the organ’s significance in adaptive immunity and as a site of metastasis. While this topic did include more traditional vaccines against different cancer types like melanoma [45], it also encompassed therapies that aimed at activating the immune system, like modulating the lymphocyte populations within the tumor [46], reprogramming the tumor microenvironment [47], or heightening the patients’ immune response [48]. CpG ODNs were commonly seen in these studies as an immune stimulator and adjuvant. Notably, the cancer vaccine field is much larger than what is represented in the current dataset because of the exclusion of vaccines in search string design.

#### Multi-modal Therapies

The Multi-modal Therapies Topic encapsulated papers that employ more than one strategy. Photothermal Therapy, Photodynamic Therapy, or magnetic fields combined with chemotherapy or immunotherapy. Interestingly, we found glioblastoma to be a prevalent cancer type in this category. These studies often utilized the coactive relationship between nanoparticles and the second strategy, like how metal nanoparticles will be rapidly heated during photothermal therapy, allowing targeted heat to the tumor and minimizing off-target effect [49]–[51]. Another second strategy seen was an alternative delivery method, like inhalation [52], [53]. As stated previously, there was overlap into the imaging category, with the second strategy being the imaging component, assisted by the nanoparticles. Some of these overlapping studies included nanoparticles being used as radiosensitizers [54], where nanoparticles enhance a tumor’s sensitivity to radiation through ROS production [55]. Overall, this topic’s studies focused on how nanoparticles and a second strategy or therapy can amplify each other’s effects, resulting in a synergistic relationship.

### Method to Implement a Living Scoping Review

While the topic model provided a great assessment of the existing literature, this review will eventually become outdated and lose its utility. Thus, we sought to determine whether we could use the eLDA topic model to generate a living review, a system that could be continuously updated **(Figure 5A)** [16], [17], [56]. We used the eLDA topic model to extract topics from 95 abstracts that also passed the full-text screening. The model was able to identify five out of the six topics in this new dataset (Figure 5B). The only topic that was not extracted was the Exosomes/Gene Delivery topic.

**Figure 5:**
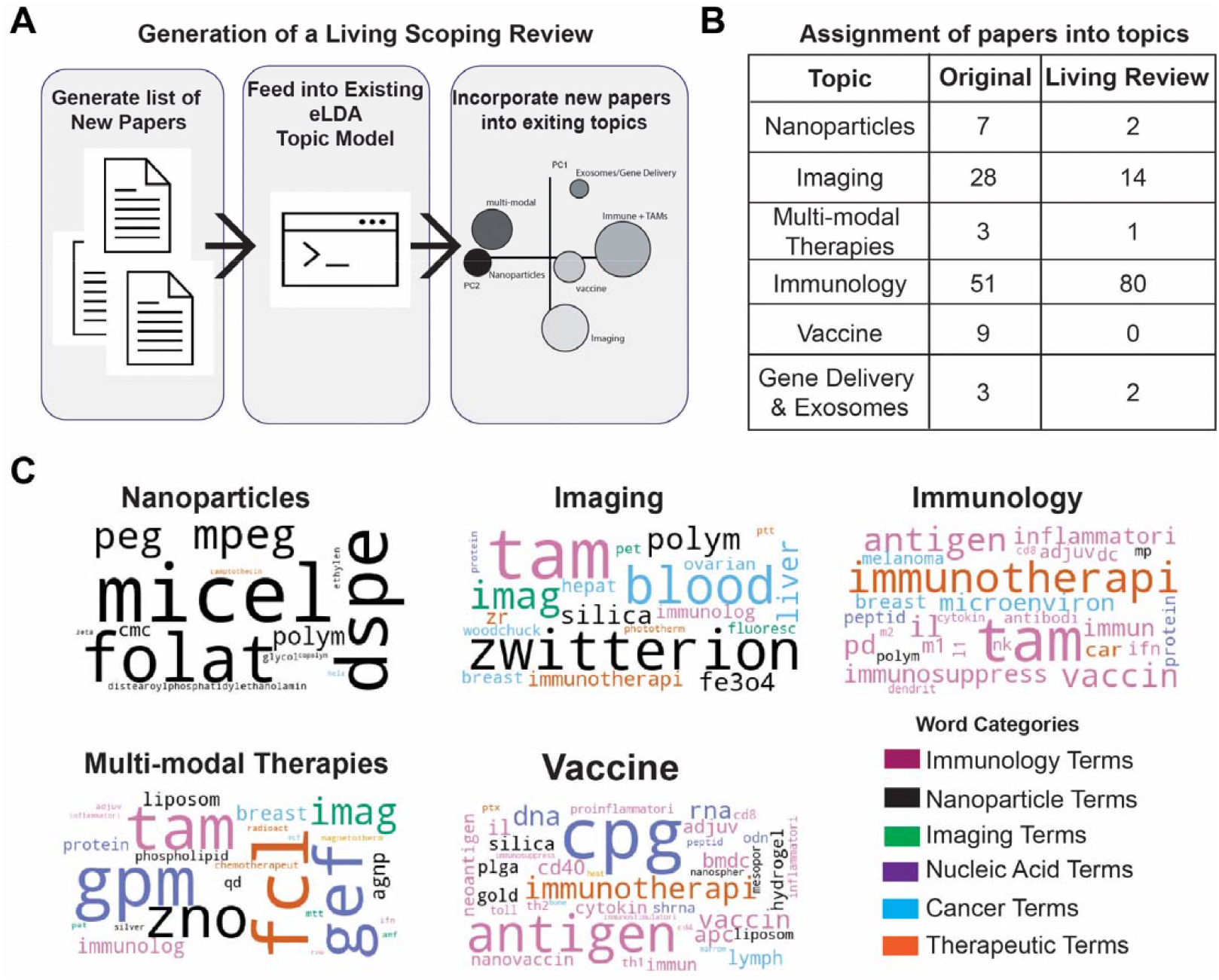
Living Review. (a) Schematic of process used to generate a living scoping review. (b) Table comparing the percentages of papers that were distributed into each topic between the original dataset and the newly acquired papers that form the Living Review. (c) Word cloud of most significant word by topic of papers included in the Living Review. Word size was determined by the frequency within the dataset while the colors were determined by their category.

This is unsurprising since this topic was the smallest in the original dataset as visualized via the intertopic distance map (Figure 3). We then followed the same methodology to identify trends within the topics using wordclouds (Figure 5C). We found there was a large degree of consistency between frequent words in the original topics and in the newer topics. For example, the Immunology topic still contained words related to TAMs and immunotherapy. Additionally, the Nanoparticles topic still had a focus on polymeric nanoparticle terms such as Micelle, DSPE, and PEG.

## Conclusion

We used machine learning enabled topic modeling to categorize pre-clinical literature regarding macrophage targeting nanoparticle literature within the cancer field. By using the eLDA model, we were able to analyze a huge subset of data collected during our scoping review. To systematically retrieve data from over 800 papers by hand is a huge undertaking and would have required an unrealistic time investment.

The eLDA topic modeling analysis revealed important insights on the last 20 years of macrophage-targeted cancer nanotherapeutics. Based on word frequency, six distinct topics were determined and labeled as “Immunology, Nanoparticles, Imaging, Gene Delivery and Exosomes, Vaccines, and Multi-modal Therapies.” The topics demonstrated different study approaches used within the field. The immunology topic highlighted the TAM repolarization strategy, where researchers attempt to make the tumor microenvironment less immunosuppressive by changing macrophage phenotype. As demonstrated in the topic’s word cloud, these papers often used liposomes and targeted breast cancer, lung cancer, or melanoma. The nanoparticle topic focused on studies whose analysis was centered on nanoparticle formulation with detailed information about the formulation’s PD/PK and structure. The nanoparticles most often studied in this fashion were polymeric and were often loaded with chemotherapeutics. The imaging topic demonstrated the strategy of using TAM’s phagocytic ability to uptake nanoparticles, often metal, to aid in imaging tumors. The most common targets for these imaging studies were shown to be tumors in the breast, liver, and brain. The exosome topic very clearly illustrated the link between using exosomes or other vesicle-like particles for gene delivery, particularly miRNA. The vaccine topic encompassed “traditional” vaccines where nanoparticles are used to deliver an antigen often alongside an adjuvant, but also studies aimed at simply activating the immune system to encourage an immune response to the tumor. CpG is very prominent within this category, expectedly, as it is a strong innate cell activator and a vaccine adjuvant. The multi-modal topic highlighted how some studies combine approaches for a synergistic treatment strategy, with NIR (near infrared radiation), photothermal therapy, and imaging being augmented by the administered nanoparticle.

While each topic employed nanoparticles, the eLDA model was able to capture distinct nanoparticle types and therapeutic features, demonstrating the scope of the TAM-targeted nanoparticle field. The word frequency tables generated using the eLDA model also allowed visualization of different cancer subtypes and organs associated with each topic as outlined above. This insight is invaluable for future therapeutic design and development, as well as an ability to draw connections between nanoparticle strategy and targeted cancer.

This study demonstrates how using Topic Modeling allows scientists to analyze large amounts of literature in a manner that can reduce prestige, journal, and geographical bias. However, as with all uses of machine learning, care must be given to the initial prompt used when generating the model to achieve the most accurate results. also recommended to collaborate with information specialists who have expertise in evidence synthesis methods when compiling search strategies to ensure comprehensive results.

It is important to consider that the topic modeling approach is a way to extract latent themes from a dataset. Data can often be summarized and categorized in multiple different ways, which is reflected in the fact that two topic models that are created using the same parameters and data can produce different results. Instead, this topic model should be interpreted as one way to organize and make sense of a large amount of information.

Finally, we demonstrated the ability to generate a living review by using our topic model to classify papers that were not included in the original dataset. Thus, we propose that topic modeling approaches can be adopted by an individual or a team of investigators for use in curating existing documents stored in reference managers. Topic modeling is a useful tool that can also be used to assess new scientific fields quickly and thoroughly.

## Supporting information

Supplementary File 1 Search Strategies

Supplementary File 2 Included Abstracts

Supplementary File 3 Code Submission

## Acknowledgements

Research reported in this publication was supported by the National Institute of General Medical Sciences of the National Institutes of Health under award numbers 1 R35 GM142957-01 and the Carnegie Mellon University SURG and ChESS Program. We would also like to thank Megha Anand, Anushree Gupta, Archippe Mbembo, Wonhee Han, Jacob Bauldock, Chris Hunyh, Hannah Yankello and Dasia Aldarondo for their participation in the abstract screening.

## Data Availability

The raw data required to reproduce these findings are included in the supplemental files. The processed data included the codes used to produce the findings are also included in the supplemental files.

## References

[1] Y. Mo, G. Kontonatsios, and S. Ananiadou, “Supporting systematic reviews using LDA-based document representations,” Syst. Rev., vol. 4, no. 1, p. 172, Nov. 2015, doi: 10.1186/s13643-015-0117-0.

[2] I. J. Marshall and B. C. Wallace, “Toward systematic review automation: a practical guide to using machine learning tools in research synthesis,” Syst. Rev., vol. 8, no. 1, p. 163, Jul. 2019, doi: 10.1186/s13643-019-1074-9.

[3] S. Bird, E. Loper, and E. Klein, “Natural Language Processing with Python,” O’Reilly Media, Inc., 2009. Accessed: May 22, 2023. [Online]. Available: https://www.nltk.org/

[4] C. B. Asmussen and C. Møller, “Smart literature review: a practical topic modelling approach to exploratory literature review,” J. Big Data, vol. 6, no. 1, p. 93, Oct. 2019, doi: 10.1186/s40537-019-0255-7.

[5] S. Pouliliou, C. Nikolaidis, and G. Drosatos, “Current trends in cancer immunotherapy: a literature-mining analysis,” Cancer Immunol. Immunother. CII, vol. 69, no. 12, pp. 2425–2439, Dec. 2020, doi: 10.1007/s00262-020-02630-8.

[6] T. Porturas and R. A. Taylor, “Forty years of emergency medicine research: Uncovering research themes and trends through topic modeling,” Am. J. Emerg. Med., vol. 45, pp. 213–220, Jul. 2021, doi: 10.1016/j.ajem.2020.08.036.

[7] R. Xu, A. Baghaei Lakeh, and N. Ghaffarzadegan, “Examining the characteristics of impactful research topics: A case of three decades of HIV-AIDS research,” J. Informetr., vol. 15, no. 1, p. 101122, Feb. 2021, doi: 10.1016/j.joi.2020.101122.

[8] D. M. Blei, A. Y. Ng, and M. I. Jordan, “Latent dirichlet allocation,” J. Mach. Learn. Res., vol. 3, no. null, pp. 993–1022, Mar. 2003.

[9] M. D. J. Peters et al., “Scoping reviews: reinforcing and advancing the methodology and application,” Syst. Rev., vol. 10, no. 1, p. 263, Oct. 2021, doi: 10.1186/s13643-021-01821-3.

[10] C. Bilynsky et al., “Scoping Review of Pre-clinical and Translational Studies on Macrophage Polarization in Nanoparticle-based Cancer Immunotherapy,” Dec. 2020, Accessed: Aug. 09, 2021. [Online]. Available: https://osf.io/pu7qb/

[11] A. Tricco et al., “PRISMA Extension for Scoping Reviews (PRISMA-ScR): Checklist and Explanation,” Ann Intern Med, vol. 169, no. 7, pp. 467–473, 2018, doi: 10.7326/M18-0850.

[12] T. J. Bozada, J. Borden, J. Workman, M. Del Cid, J. Malinowski, and T. Luechtefeld, “Sysrev: A FAIR Platform for Data Curation and Systematic Evidence Review,” Front. Artif. Intell., vol. 0, 2021, doi: 10.3389/frai.2021.685298.

[13] M. Röder, A. Both, and A. Hinneburg, “Exploring the Space of Topic Coherence Measures,” in Proceedings of the Eighth ACM International Conference on Web Search and Data Mining, in WSDM ‘15. New York, NY, USA: Association for Computing Machinery, Feb. 2015, pp. 399–408. doi: 10.1145/2684822.2685324.

[14] “Ensemble LDA | sezanzeb.de.” https://www.sezanzeb.de/machine_learning/ensemble_LDA/ (accessed Jun. 08, 2023).

[15] R. Řehůřek and P. Sojka, Software Framework for Topic Modelling with Large Corpora. 2010, p. 50. doi: 10.13140/2.1.2393.1847.

[16] M. Kelechava, “Using LDA Topic Models as a Classification Model Input,” Towards Data Science, Mar. 03, 2019. https://towardsdatascience.com/unsupervised-nlp-topic-models-as-a-supervised-learning-input-cf8ee9e5cf28 (accessed Jun. 08, 2023).

[17] H. B. Yalamanchili, S. J. Kho, and M. L. Raymer, “Latent Dirichlet Allocation for Classification using Gene Expression Data,” in 2017 IEEE 17th International Conference on Bioinformatics and Bioengineering (BIBE), Oct. 2017, pp. 39–44. doi: 10.1109/BIBE.2017.00-81.

[18] M. J. Page et al., “The PRISMA 2020 statement: an updated guideline for reporting systematic reviews,” BMJ, vol. 372, p. n71, Mar. 2021, doi: 10.1136/bmj.n71.

[19] J. B. Lee, H. R. Kim, and S.-J. Ha, “Immune Checkpoint Inhibitors in 10 Years: Contribution of Basic Research and Clinical Application in Cancer Immunotherapy,” Immune Netw., vol. 22, no. 1, Feb. 2022, doi: 10.4110/in.2022.22.e2.

[20] D. Hanahan and R. A. Weinberg, “Hallmarks of cancer: the next generation,” Cell, vol. 144, no. 5, pp. 646–674, Mar. 2011, doi: 10.1016/j.cell.2011.02.013.

[21] “First-Ever CAR T-cell Therapy Approved in U.S,” Cancer Discov., vol. 7, no. 10, p. OF1, Oct. 2017, doi: 10.1158/2159-8290.CD-NB2017-126.

[22] F. J. van Dalen, M. H. M. E. van Stevendaal, F. L. Fennemann, M. Verdoes, and O. Ilina, “Molecular Repolarisation of Tumour-Associated Macrophages,” Molecules, vol. 24, no. 1, p. 9, Dec. 2018, doi: 10.3390/molecules24010009.

[23] D. Mimno, H. Wallach, E. Talley, M. Leenders, and A. McCallum, “Optimizing Semantic Coherence in Topic Models,” in Proceedings of the 2011 Conference on Empirical Methods in Natural Language Processing, Edinburgh, Scotland, UK.: Association for Computational Linguistics, Jul. 2011, pp. 262–272. Accessed: May 22, 2023. [Online]. Available: https://aclanthology.org/D11-1024

[24] “Welcome to pyLDAvis’s documentation! — pyLDAvis 2.1.2 documentation.” https://pyldavis.readthedocs.io/en/latest/ (accessed May 22, 2023).

[25] D.-Y. Gao et al., “CXCR4-targeted lipid-coated PLGA nanoparticles deliver sorafenib and overcome acquired drug resistance in liver cancer,” Biomaterials, vol. 67, pp. 194–203, Oct. 2015, doi: 10.1016/j.biomaterials.2015.07.035.

[26] R. Arakaki et al., “CCL2 as a potential therapeutic target for clear cell renal cell carcinoma,” Cancer Med., vol. 5, no. 10, pp. 2920–2933, Sep. 2016, doi: 10.1002/cam4.886.

[27] Y. Zhang et al., “Glycocalyx-Mimicking Nanoparticles Improve Anti-PD-L1 Cancer Immunotherapy through Reversion of Tumor-Associated Macrophages,” Biomacromolecules, vol. 19, no. 6, pp. 2098–2108, Jun. 2018, doi: 10.1021/acs.biomac.8b00305.

[28] Y. L. Li, Y. T. Chen, N. V. Cuong, and M. F. Hsieh, “Preparation of Nanoparticles of AB2 Triblock Copolymers for Doxorubicin Delivery,” in 6th World Congress of Biomechanics (WCB 2010). August 1-6, 2010 Singapore, C. T. Lim and J. C. H. Goh, Eds., in IFMBE Proceedings. Berlin, Heidelberg: Springer, 2010, pp. 1242–1245. doi: 10.1007/978-3-642-14515-5_315.

[29] X. Que et al., “Study on preparation, characterization and multidrug resistance reversal of red blood cell membrane-camouflaged tetrandrine-loaded PLGA nanoparticles,” Drug Deliv., vol. 26, no. 1, pp. 199–207, Dec. 2019, doi: 10.1080/10717544.2019.1573861.

[30] N.-V. Cuong, Y.-T. Chen, and M.-F. Hsieh, “Doxorubicin-loaded micelles of y-shaped peg-(pcl)2 against drug-resistant breast cancer cells,” Biomed. Eng. Appl. Basis Commun., vol. 25, no. 05, p. 1340009, Oct. 2013, doi: 10.4015/S1016237213400097.

[31] C. Castellani et al., “Tumor-facing hepatocytes significantly contribute to mild hyperthermia-induced targeting of rat liver metastasis by PLGA-NPs,” Int. J. Pharm., vol. 566, pp. 541–548, Jul. 2019, doi: 10.1016/j.ijpharm.2019.06.004.

[32] Y. Sui et al., “Tumor-specific design of PEGylated gadolinium-based nanoscale particles: Facile synthesis, characterization, and improved magnetic resonance imaging of metastasis lung cancer,” J. Photochem. Photobiol. B, vol. 202, p. 111669, Jan. 2020, doi: 10.1016/j.jphotobiol.2019.111669.

[33] Z. Zhou et al., “A Protein-Corona-Free T1–T2 Dual-Modal Contrast Agent for Accurate Imaging of Lymphatic Tumor Metastasis,” ACS Appl. Mater. Interfaces, vol. 7, no. 51, pp. 28286–28293, Dec. 2015, doi: 10.1021/acsami.5b08422.

[34] H. Vu-Quang et al., “Targeted delivery of mannan-coated superparamagnetic iron oxide nanoparticles to antigen-presenting cells for magnetic resonance-based diagnosis of metastatic lymph nodes in vivo,” Acta Biomater., vol. 7, no. 11, pp. 3935–3945, Nov. 2011, doi: 10.1016/j.actbio.2011.06.044.

[35] Y. Li et al., “Targeted Imaging of CD206 Expressing Tumor-Associated M2-like Macrophages Using Mannose-Conjugated Antibiofouling Magnetic Iron Oxide Nanoparticles,” ACS Appl. Bio Mater., vol. 3, no. 7, pp. 4335–4347, Jul. 2020, doi: 10.1021/acsabm.0c00368.

[36] A. Leimgruber et al., “Behavior of endogenous tumor-associated macrophages assessed in vivo using a functionalized nanoparticle,” Neoplasia N. Y. N, vol. 11, no. 5, pp. 459–468, 2 p following 468, May 2009, doi: 10.1593/neo.09356.

[37] L. D. Tran et al., “Nanosized magnetofluorescent Fe3O4–curcumin conjugate for multimodal monitoring and drug targeting,” Colloids Surf. Physicochem. Eng. Asp., vol. 371, no. 1–3, pp. 104–112, Nov. 2010, doi: 10.1016/j.colsurfa.2010.09.011.

[38] J. Janjic, S. K. Patel, M. Patrick, E. Divito, and M. Cascio, Suppressing inflammation from inside out with novel NIR visible perfluorocarbon nanotheranostics, vol. 8596. 2013, p. 85960L. doi: 10.1117/12.2004625.

[39] R. Kalluri and V. S. LeBleu, “The biology, function, and biomedical applications of exosomes,” Science, vol. 367, no. 6478, p. eaau6977, Feb. 2020, doi: 10.1126/science.aau6977.

[40] P. Jiang et al., “CD137 promotes bone metastasis of breast cancer by enhancing the migration and osteoclast differentiation of monocytes/macrophages,” Theranostics, vol. 9, no. 10, pp. 2950–2966, 2019, doi: 10.7150/thno.29617.

[41] E. Cho et al., “Comparison of exosomes and ferritin protein nanocages for the delivery of membrane protein therapeutics,” J. Control. Release Off. J. Control. Release Soc., vol. 279, pp. 326–335, Jun. 2018, doi: 10.1016/j.jconrel.2018.04.037.

[42] S. Bhattacharyya and S. S. Ghosh, “Transmembrane TNFα-Expressed Macrophage Membrane-Coated Chitosan Nanoparticles as Cancer Therapeutics,” ACS Omega, vol. 5, no. 3, pp. 1572–1580, Jan. 2020, doi: 10.1021/acsomega.9b03531.

[43] J. Zhu et al., “A Biohybrid Lurker-to-Attacker Strategy To Solve Inherent Dilemma of Positively Charged Delivery Nanoparticles,” Chem. Mater., vol. 29, no. 5, pp. 2227–2231, Mar. 2017, doi: 10.1021/acs.chemmater.6b05120.

[44] P. L. Rodriguez, T. Harada, D. A. Christian, D. A. Pantano, R. K. Tsai, and D. E. Discher, “Minimal ‘Self’ Peptides That Inhibit Phagocytic Clearance and Enhance Delivery of Nanoparticles,” Science, vol. 339, no. 6122, pp. 971–975, Feb. 2013, doi: 10.1126/science.1229568.

[45] G. Zhu et al., “DNA-inorganic hybrid nanovaccine for cancer immunotherapy,” Nanoscale, vol. 8, no. 12, pp. 6684–6692, Mar. 2016, doi: 10.1039/c5nr08821f.

[46] D. H. Kim et al., “Liposome-encapsulated CpG enhances antitumor activity accompanying the changing of lymphocyte populations in tumor via intratumoral administration,” Nucleic Acid Ther., vol. 25, no. 2, pp. 95–102, Apr. 2015, doi: 10.1089/nat.2014.0509.

[47] L. Munakata et al., “Lipid nanoparticles of Type-A CpG D35 suppress tumor growth by changing tumor immune-microenvironment and activate CD8 T cells in mice,” J. Control. Release Off. J. Control. Release Soc., vol. 313, pp. 106–119, Nov. 2019, doi: 10.1016/j.jconrel.2019.09.011.

[48] J. Meng et al., “Subcutaneous injection of water-soluble multi-walled carbon nanotubes in tumor-bearing mice boosts the host immune activity,” Nanotechnology, vol. 21, no. 14, p. 145104, Mar. 2010, doi: 10.1088/0957-4484/21/14/145104.

[49] S. J. Madsen et al., “Nanoparticle-loaded macrophage-mediated photothermal therapy: potential for glioma treatment,” Lasers Med. Sci., vol. 30, no. 4, pp. 1357–1365, May 2015, doi: 10.1007/s10103-015-1742-5.

[50] Y. Gao et al., “Targeted photothermal therapy of mice and rabbits realized by macrophage-loaded tungsten carbide,” Biomater. Sci., vol. 7, no. 12, pp. 5350–5358, Nov. 2019, doi: 10.1039/C9BM00911F.

[51] B. Ji et al., “Hybrid membrane camouflaged copper sulfide nanoparticles for photothermal-chemotherapy of hepatocellular carcinoma,” Acta Biomater., vol. 111, pp. 363–372, Jul. 2020, doi: 10.1016/j.actbio.2020.04.046.

[52] Y. Yang, N. Bajaj, P. Xu, K. Ohn, M. D. Tsifansky, and Y. Yeo, “Development of highly porous large PLGA microparticles for pulmonary drug delivery,” Biomaterials, vol. 30, no. 10, pp. 1947–1953, Apr. 2009, doi: 10.1016/j.biomaterials.2008.12.044.

[53] L. Zhu, M. Li, X. Liu, L. Du, and Y. Jin, “Inhalable oridonin-loaded poly(lactic-co-glycolic)acid large porous microparticles for in situ treatment of primary non-small cell lung cancer,” Acta Pharm. Sin. B, vol. 7, no. 1, pp. 80–90, Jan. 2017, doi: 10.1016/j.apsb.2016.09.006.

[54] M. Enferadi et al., “Radiosensitization of ultrasmall GNP-PEG-cRGDfK in ALTS1C1 exposed to therapeutic protons and kilovoltage and megavoltage photons,” Int. J. Radiat. Biol., vol. 94, no. 2, pp. 124–136, Feb. 2018, doi: 10.1080/09553002.2018.1407462.

[55] C. Bilynsky, N. Millot, and A.-L. Papa, “Radiation nanosensitizers in cancer therapy—From preclinical discoveries to the outcomes of early clinical trials,” Bioeng. Transl. Med., vol. n/a, no. n/a, p. e10256, doi: 10.1002/btm2.10256.

[56] J. H. Elliott et al., “Living systematic review: 1. Introduction—the why, what, when, and how,” J. Clin. Epidemiol., vol. 91, pp. 23–30, Nov. 2017, doi: 10.1016/j.jclinepi.2017.08.010.

