## Supplementary File 1 Search Strategies for "Meta-analysis of macrophage nanoparticle targeting across blood and solid tumors using an eLDA Topic modeling Machine Learning approach"

**Supplemental File: Search strategies used to find relevant studies for screening**

**PubMed (Medline)**

| 1 | Nanoparticles[MeSH] |
| --- | --- |
| 2 | Nanoparticle*[tiab] OR dendrimer*[tiab] OR nanocapsule*[tiab] OR nanoconjugate*[tiab] OR nanosphere*[tiab] OR nanotechnolog*[tiab] OR nanogel*[tiab] OR nanomedicine*[tiab] OR nanorod*[tiab] OR nanotube*[tiab] OR liposom*[tiab] OR quantum dot*[tiab] OR polymer[tiab] OR polymers[tiab] OR polymeric[tiab] |
| 3 | 1 OR 2 |
| 4 | Macrophages[MeSH] OR monocytes[Mesh] |
| 5 | Macrophage*[tiab] OR monocyt*[tiab] |
| 6 | Microglia*[tiab] OR Kupffer*[tiab] OR Langerhans[tiab] OR TAMS[tiab] |
| 7 | 4 OR 5 OR 6 |
| 8 | Neoplasms[MeSH] OR Immunotherapy[MeSH] OR drug therapy[MeSH] |
| 9 | Cancer*[tiab] OR tumor*[tiab] OR tumour*[tiab] OR immunotherap*[tiab] OR chemotherap*[tiab] OR immunomodulatory[tiab] OR immune[tiab] OR immuni*[tiab] OR carcinoma*[tiab] OR sarcoma*[tiab] OR lymphoma*[tiab] OR leukemia[tiab] OR myeloma[tiab] OR malignan*[tiab] OR neoplasm*[tiab] OR drug therapy[tiab] OR drug therapies[tiab] OR drug delivery[tiab] |
| 10 | 8 OR 9 |
| 11 | 3 AND 7 AND 10 |
| 12 | Editorial[pt] OR Letter[pt] OR "Newspaper Article"[pt] OR Comment[pt] |
| 13 | #11 NOT #12 |
| 14 | Limit to English |
| 15 | Limit Date 2000-2020 |

**Web of Science Core Collection (Clarivate) including: Science Citation Index-Expanded (SCI-Expanded), Social Science Citation Index (SSCI) and Emerging Sources Citation Index (ESCI)**

| 1 | TS=(Nanoparticle* OR dendrimer* OR nanocapsule* OR nanoconjugate* OR nanosphere* OR nanotechnolog* OR nanogel* OR nanomedicine* OR nanorod* OR nanotube* OR liposom* OR "quantum dot*" OR polymer OR polymers OR polymeric) |
| --- | --- |
| 2 | TS=(Macrophage* OR monocyt*) |
| 3 | TS=(Microglia* OR Kupffer* OR Langerhans OR TAMS) |
| 4 | 2 OR 3 |
| 5 | TS=(Cancer* OR tumor* OR tumour* OR immunotherap* OR chemotherap* OR immunomodulatory OR immune OR immuni* OR carcinoma* OR sarcoma* OR lymphoma* OR leukemia OR myeloma[tiab] OR malignan* OR neoplasm* OR "drug therapy" OR "drug therapies" OR "drug delivery") |
| 6 | 1 AND 4 AND 5 |
| 7 | Exclude Document Types: Book Chapter, Meeting Abstract, Editorial Material, News Item, Letter |
| 9 | Limit Date 2000-2020 |

**Scopus (Elsevier)**

| 1 | TITLE-ABS-KEY(Nanoparticle* OR dendrimer* OR nanocapsule* OR nanoconjugate* OR nanosphere* OR nanotechnolog* OR nanogel* OR nanomedicine* OR nanorod* OR nanotube* OR liposom* OR "quantum dot*" OR polymer OR polymers OR polymeric) |
| --- | --- |
| 2 | TITLE-ABS-KEY(Macrophage* OR monocyt*) |
| 3 | TITLE-ABS-KEY(Microglia* OR Kupffer* OR Langerhans OR TAMS) |
| 4 | 2 OR 3 |
| 5 | TITLE-ABS-KEY(Cancer* OR tumor* OR tumour* OR immunotherap* OR chemotherap* OR immunomodulatory OR immune OR immuni* OR carcinoma* OR sarcoma* OR lymphoma* OR leukemia OR myeloma OR malignan* OR neoplasm* OR "drug therapy" OR "drug therapies" OR "drug delivery") |
| 6 | 1 AND 4 AND 5 |
| 7 | LIMIT-TO (LANGUAGE , "English")) AND (LIMIT-TO (DOCTYPE, "ar", "re", "cp" ) |
| 8 | Limit Date 2000-2020 |

**IEEE Xplore**

("All Metadata":Nanoparticle OR nanoparticles OR dendrimer OR dendrimers OR nanocapsule* OR nanoconjugate* OR nanosphere* OR nanotechnology OR nanogel OR nanogels OR nanomedicine OR nanomedicines OR nanorod OR nanorods OR nanotube* OR liposome OR liposomes OR "quantum dot" OR "quantum dots" OR polymer OR polymers OR polymeric) AND ("All Metadata":Macrophage OR macrophages OR monocyte OR monocytic OR monocytes OR Microglia OR Kupffer OR Langerhans OR TAMS)

Limit Date 2000-2020, Limit Language to English

**Biotechnology & BioEngineering Abstracts (ProQuest)**

| 1 | TI(Nanoparticle* OR dendrimer* OR nanocapsule* OR nanoconjugate* OR nanosphere* OR nanotechnolog* OR nanogel* OR nanomedicine* OR nanorod* OR nanotube* OR liposom* OR "quantum dot*" OR polymer OR polymers OR polymeric) OR AB(Nanoparticle* OR dendrimer* OR nanocapsule* OR nanoconjugate* OR nanosphere* OR nanotechnolog* OR nanogel* OR nanomedicine* OR nanorod* OR nanotube* OR liposom* OR "quantum dot*" OR polymer OR polymers OR polymeric) OR MAINSUBJECT(Nanoparticle* OR dendrimer* OR nanocapsule* OR nanoconjugate* OR nanosphere* OR nanotechnolog* OR nanogel* OR nanomedicine* OR nanorod* OR nanotube* OR liposom* OR "quantum dot*" OR polymer OR polymers OR polymeric) |
| --- | --- |
| 2 | TI(Macrophage* OR monocyt*) OR AB(Macrophage* OR monocyt*) OR MAINSUBJECT(Macrophage* OR monocyt*) |
| 3 | TI(Microglia* OR Kupffer* OR Langerhans OR TAMS) OR AB(Microglia* OR Kupffer* OR Langerhans OR TAMS) OR MAINSUBJECT(Microglia* OR Kupffer* OR Langerhans OR TAMS) |
| 4 | 2 OR 3 |
| 5 | TI(Cancer* OR tumor* OR tumour* OR immunotherap* OR chemotherap* OR immunomodulatory OR immune OR immuni* OR carcinoma* OR sarcoma* OR lymphoma* OR leukemia OR myeloma[tiab] OR malignan* OR neoplasm* OR "drug therapy" OR "drug therapies" OR "drug delivery") OR AB(Cancer* OR tumor* OR tumour* OR immunotherap* OR chemotherap* OR immunomodulatory OR immune OR immuni* OR carcinoma* OR sarcoma* OR lymphoma* OR leukemia OR myeloma[tiab] OR malignan* OR neoplasm* OR "drug therapy" OR "drug therapies" OR "drug delivery") OR MAINSUBJECT(Cancer* OR tumor* OR tumour* OR immunotherap* OR chemotherap* OR immunomodulatory OR immune OR immuni* OR carcinoma* OR sarcoma* OR lymphoma* OR leukemia OR myeloma[tiab] OR malignan* OR neoplasm* OR "drug therapy" OR "drug therapies" OR "drug delivery") |
| 6 | 1 AND 4 AND 5 |
| 7 | Exclude Source Type: Book, Trade Journals |
| 8 | Limit Date 2000-2020 |

**Google Scholar**

nanoparticle AND (macrophage OR monocyte) AND (cancer OR chemotherapy OR malignant OR malignancy OR neoplasm)

First 1000 results scraped with Data Miner (<https://data-miner.io/>) and deduplicated against previous searches.
