## Supplementary File 3 Code Submission for "Meta-analysis of macrophage nanoparticle targeting across blood and solid tumors using an eLDA Topic modeling Machine Learning approach"

*Objective: The purpose of this notebook is to provide example code that was used for the creation of the paper 'Overview of macrophage targeting across blood and solid tuors using an eLDA Topic Modeling Machine Learning Approach*

Sources:

- LDA Example Code: https://rstudio-pubs-static.s3.amazonaws.com/79360_850b2a69980c4488b1db95987a24867a.html*
- Pandas Documentation: https://pandas.pydata.org/docs/reference/api/pandas.DataFrame.html?highlight=dataframe#pandas.DataFrame
- NLTK RegExp Documentation: http://www.nltk.org/api/nltk.tokenize.html?highlight=regexp
- Info on Python Regular Expressions: https://docs.python.org/3/library/re.html
- Citation to paper that also used common stopwords: https://www.sciencedirect.com/science/article/abs/pii/S0735675720307221?via%3Dihub (This is citation 5 in the paper)
- Stemmer Documentation: https://www.nltk.org/api/nltk.stem.html.
- Porter Stemer Algorithm webpage: https://tartarus.org/martin/PorterStemmer/
- Gensim: https://radimrehurek.com/gensim/intro.html#:~:text=Gensim%20is%20a%20free%20open,human%2Dwise)%20as%20possible.
- Word Cloud Code: https://amueller.github.io/word_cloud/auto_examples/colored_by_group.html
- Towards Data Science LDA Classifier Article: https://towardsdatascience.com/unsupervised-nlp-topic-models-as-a-supervised-learning-input-cf8ee9e5cf28

#### Step 1: Importing Documents

This is where we will import the data sets. There are two important datasets we will be importing to this file. The file (df_docs) contains the documents that will be used to train the model.

There are 854 total documents in the dataset. We will specifically be using the Abstract column for this model. However, the other columns will become important for the purpose of analysis later on.

```
# mount Google Drive
from google.colab import drive
drive.mount('/content/grive')
```

```
Mounted at /content/grive
```

```
import pandas as pd

# Define file path
path = ''
# Import data frame
df = pd.read_csv(path)

# Get abstracts
document_set = df['Abstract']
print('Abstracts:')
print(document_set)

# Get article ID's
article_id = df['Article ID']
print('\nArticle ID:')
print(article_id)

# Compile features of dcument set
N_docs = len(document_set)

print("\nNumber of documents: %d documents" %N_docs)
```

```
Abstracts:
0      Biomimetic nanoengineering built through integ...
1      Zoledronic acid (ZOL) has been used as an adju...
2      Triple negative breast cancer (TNBC) is the de...
3      Lung cancer remains the leading cause of cance...
4      The thermoplasmonic properties of platinum nan...
                             ...
849    Cyclic dinucleotides are of importance in the ...
850    Advances in genetic engineering tools have con...
851    To engineer patient-derived cells into therapy...
852    Synergistic phototherapy has the potential to ...
853    There is a tremendous focus on the application...
Name: Abstract, Length: 854, dtype: object

Article ID:
0       8334487
1       8334505
2       8334684
3       8334718
4       8334819
         ...
849    10422145
850    10422183
851    10422186
852    10422188
853    10422210
Name: Article ID, Length: 854, dtype: int64

Number of documents: 854 documents
```

#### Step 2 - Cleaning Documents

Before applying any machine algorithm, the data needs to be cleaned. There are three steps in data cleaning:

1. Tokenization
2. Removing Stop Words
3. Stemming

##### Step 2.1 - Tokenization

First, we need to separate the document into its basic elements:words.

Tokenization will be done using the `tokenize.regexp` module. This tokenizer takes in a regular expression to determine how to tokenize the word. The output is a list of tokenized words from the dictionary.

This tokenizer will only recognize words in the English alphabet (a-z). It will not recognize numbers, contractions, possessivee nouns, etc.

- The `r` preceeding the regex string removes the need for an additional backslash to escape the first one for every special character. This is called Raw String Notation.
- `[\w]` + matches Unicode word characters

The try-except statement is there to handle the case that the abstract is empty. Otherwise, there would be an Attribute Error from trying to convert a float (nan) to a lowercase string.

```
from nltk.tokenize import RegexpTokenizer

# Create the tokenizer
# Regex matches to Unicode word characters
tokenizer = RegexpTokenizer(r'\w+')


# Define function that tokenizes a string
def tokenize_document(document):
  try:
    # Convert document to lowercase
    raw_document = document.lower()

    # Tokenize the raw document and add it to tokens
    tokenized_document = tokenizer.tokenize(raw_document)

    # Return the tokenized document
    return tokenized_document

  # Handle case of empty abstract
  except AttributeError:
    return []
```

```
# Apply tokenizer to abstracts
tokens = document_set.apply(tokenize_document)

# Print number of tokenized documents
print("Number of tokenized documents: %d" %len(tokens))

print("\nFirst 10 tokenized documents:")
for i in range(10):
  print("\nDocument %d:" %article_id[i])
  print(tokens[i])
```

```
Number of tokenized documents: 854

First 10 tokenized documents:

Document 8334487:
['biomimetic', 'nanoengineering', 'built', 'through', 'integrating', 'the', '

Document 8334505:
['zoledronic', 'acid', 'zol', 'has', 'been', 'used', 'as', 'an', 'adjuvant',

Document 8334684:
['triple', 'negative', 'breast', 'cancer', 'tnbc', 'is', 'the', 'deadliest',

Document 8334718:
['lung', 'cancer', 'remains', 'the', 'leading', 'cause', 'of', 'cancer', 'rel

Document 8334819:
['the', 'thermoplasmonic', 'properties', 'of', 'platinum', 'nanoparticles', '

Document 8335232:
['the', 'pursuit', 'of', 'more', 'selectivity', 'in', 'the', 'delivery', 'of'

Document 8335457:
['patients', 'with', 'malignant', 'ascites', 'mas', 'display', 'several', 'sy

Document 8335543:
['iron', 'oxide', 'nanoparticles', 'have', 'found', 'widespread', 'applicatio

Document 8335571:
['previous', 'studies', 'have', 'identified', 'relevant', 'genes', 'and', 'si

Document 8336594:
['a', 'kind', 'of', 'x', 'ray', 'computed', 'tomography', 'ct', 'fluorescence
```

##### Step 2.2 - Removing Stop Words

Certain words, such as 'the', 'I', and 'and' are meaningless in a topic model. These words are called stop words, and they need to be removed from the text before applying the LDA model. The list of stopwords for this model was derived in three steps.

1. A list of common stopwords from NLTK
2. A list of stopwords curated from the search string used to locate records in the databases.
3. Stop words that occured in only one document or in greater than 90% of documents.
4. Words that contain numbers.

```
import nltk
from nltk.corpus import stopwords
nltk.download('stopwords')

stop_words_list = stopwords.words('english')

stop_words_list
```

```
[nltk_data] Downloading package stopwords to /root/nltk_data...
[nltk_data]   Unzipping corpora/stopwords.zip.
['i',
 'me',
 'my',
 'myself',
 'we',
 'our',
 'ours',
 'ourselves',
 'you',
 "you're",
 "you've",
 "you'll",
 "you'd",
 'your',
 'yours',
 'yourself',
 'yourselves',
 'he',
 'him',
 'his',
 'himself',
 'she',
 "she's",
 'her',
```

```
      'hers',
      'herself',
      'it',
      "it's",
      'its',
      'itself',
      'they',
      'them',
      'their',
      'theirs',
      'themselves',
      'what',
      'which',
      'who',
      'whom',
      'this',
      'that',
      "that'll",
      'these',
      'those',
      'am',
      'is',
      'are',
      'was',
      'were',
      'be',
      'been',
      'being',
      'have',
      'has',
      'had',
      'having',
      'do',
      [illegible]
```

```
# add additional stop words
nano_stop_words = ['nanoparticle', 'nanoparticles', 'nano', 'particle', 'particles'
                   'np', 'nps']
stop_words_list.extend(nano_stop_words)

macrophage_stop_words = ['macrophage', 'macrophages']
stop_words_list.extend(macrophage_stop_words)

cancer_stop_words = ['tumor', 'tumors', 'cancer', 'cancers', 'cancerous']
stop_words_list.extend(cancer_stop_words)

medical_stop_words = ['therapy', 'therapies', 'drug', 'drugs', 'cell', 'cells']
```

```
stop_words_list.extend(medical_stop_words)

other_stop_words = ['study', 'studies', 'background', 'use', 'uses', 'used',
                    'user', 'using', 'effect', 'effects', 'effective',
                    'result', 'results', 'resulted', 'can', 'abstract', 'could',
                    'uptake', 'uptakes', 'uptook', 'uptaking', 'treatment',
                    'treat', 'treats', 'treated', 'treating', 'treatments',
                    'show', 'shows', 'shown', 'showed', 'showing', 'target', 'targe
                    'targeted', 'tarteting', 'immune', 'high', 'higher', 'highest',
                    'highly', 'low', 'lower', 'lowest', 'increase', 'increased',
                    'increasing', 'increases', 'decrease', 'decreases', 'decreased'
                    'decreasing', 'might', 'may', 'cause', 'causes', 'caused', 'cau
                    'causing', 'because', 'investigate', 'investigates', 'investiga
                    'investigated', 'study', 'studies', 'studied', 'studying', 'sys
                    'systems', 'load', 'loaded', 'loads', 'loading', 'develop', 'de
                    'developing', 'vivo', 'vitro', 'target', 'targets', 'targeted',
                    'targeting', 'therapy', 'therapeutic', 'therepeutics', 'like',
                    'likes', 'liking', 'liked', 'active', 'activate', 'activated',
                    'activating', 'activates', 'deliver', 'delivery', 'deliveries',
                    'delivered', 'delivering']
stop_words_list.extend(other_stop_words)

letters = ['a', 'b', 'c', 'd', 'e', 'f', 'g', 'h', 'i', 'j', 'k', 'l', 'm', 'n',
           'o', 'p', 'q', 'r', 's', 't', 'u', 'v', 'w', 'x', 'y', 'z']
stop_words_list.extend(letters)

# print list of stop words
print("Stop words list:\n")

for word in stop_words_list:
  print(word)

print("\nNumber of stop words: %d words" %len(stop_words_list))
```

```
Stop words list:

i
me
my
myself
we
our
ours
ourselves
you
you're
you've
```

```
you've
you'll
you'd
your
yours
yourself
yourselves
he
him
his
himself
she
she's
her
hers
herself
it
it's
its
itself
they
them
their
theirs
themselves
what
which
who
whom
this
that
that'll
these
those
am
is
are
was
were
be
been
being
have
has
had
having
do
[illegible]
```

```
# remove stop words that occur once,  in <1 document or > 90% of all documents
doc_freq = {}
term_freq = {}

for doc in (tokens):
  seen = []

  for w in doc:
    term_freq[w] = term_freq.get(w, 0) + 1
    if w not in seen:
      seen.append(w)
      doc_freq[w] = doc_freq.get(w, 0) + 1


# make new list of stop words
term_threshold = 1
doc_lower_threshold = 1
doc_upper_threshold = 0.9 * N_docs

# create list of excluded terms
term_stopwords = []
term_keys = term_freq.keys()
for w in term_keys:
  if (term_freq.get(w, 0) <= term_threshold):
    term_stopwords.append(w)
stop_words_list.extend(term_stopwords)

# excluded terms based on document lowerthreshold
doc_stopwords = []
doc_keys = doc_freq.keys()
for w in doc_keys:
  if (doc_freq.get(w, 0) <= doc_lower_threshold):
    doc_stopwords.append(w)
  if (doc_freq.get(w, 0) > doc_upper_threshold):
    doc_stopwords.append(w)
stop_words_list.extend(doc_stopwords)

print('Number of stop words: ', len(stop_words_list))
```

```
    Number of stop words:  8708
```

```
# define a function that removes the stop words from a tokenized document
def stop_document(doc):

  # create a new list for that document
  stopped_doc = []

  # look at every token in the document
  for token in doc:

    # check to see if the word is a stop word
    if (not(token.isdecimal()) and not(token in stop_words_list)):

      # add token to the stopped token list
      stopped_doc.append(token)

  # return the stopped document
  return stopped_doc
```

```
# Apply stop words remover to tokenized abstracts
stopped_tokens = tokens.apply(stop_document)
```

```
# Print number of tokenized documents
print("Number of stopped documents: %d" %len(stopped_tokens))

print("\nFirst 10 stopped documents:")
for i in range(10):
  print("\nDocument %d:" %article_id[i])
  print(stopped_tokens[i])
```

```
Number of stopped documents: 854

First 10 stopped documents:

Document 8334487:
['biomimetic', 'built', 'integrating', 'specific', 'membrane', 'artificially',

Document 8334505:
['zoledronic', 'acid', 'zol', 'adjuvant', 'breast', 'suggested', 'zol', 'asso

Document 8334684:
['triple', 'negative', 'breast', 'tnbc', 'deadliest', 'form', 'breast', 'succe

Document 8334718:
['lung', 'remains', 'leading', 'related', 'death', 'united', 'states', 'althou

Document 8334819:
['properties', 'platinum', 'render', 'desirable', 'diagnosis', 'detection', '

Document 8335232:
['selectivity', 'plasmonic', 'critical', 'penetration', 'clinical', 'applicat

Document 8335457:
['patients', 'malignant', 'ascites', 'mas', 'display', 'several', 'symptoms',

Document 8335543:
['iron', 'oxide', 'found', 'widespread', 'applications', 'different', 'areas',

Document 8335571:
['previous', 'identified', 'relevant', 'genes', 'signalling', 'pathways', 'ha

Document 8336594:
['kind', 'ray', 'computed', 'tomography', 'ct', 'fluorescence', 'dual', 'moda
```

```
stop_words_df = pd.DataFrame(stop_words_list, columns = ["Words"])
stop_words_df.to_csv('/content/grive/MyDrive/ML Literature Review/Chloe Brown/Revis
```

##### Step 2.3 - Stemming

We will be using the PorterStemmer class from nltk. Documentation: https://www.nltk.org/api/nltk.stem.html.

Algorithm webpage: https://tartarus.org/martin/PorterStemmer/

For the best results, use the default parameters for the constructor when creating the stemmer (`PorterStemmer()` constructor).

We will be using the PorterStemmer class from nltk. Stemming is important becuase it removes prefixes of words. This is important because it allows for words with the same stem but different affixes to be treated the same by the LDA algorithm. For example, macrophages and macrophage should be treated as the same word by the algorithm because they refer to the same concept.

```
from nltk.stem.porter import PorterStemmer

# construct the stemmer
stemmer = PorterStemmer()


# define a function that stems words in a tokenized document
def stem_document(doc):

  # create a new list for that document
  stemmed_doc = []

  # look at every token in the document
  for token in doc:

    # stem the token and add it to the list
    stemmed_doc.append(stemmer.stem(token))

  # add newly stemmed document to the list of stememd tokens
  return stemmed_doc
```

```
# apply stemmer to dataframe
text = stopped_tokens.apply(stem_document)

# Print number of tokenized documents
print("Number of stemmed documents: %d" %len(text))

print("\nFirst 10 stemmed documents:")
for i in range(10):
  print("\nDocument %d:" %article_id[i])
  print(text[i])
```

```
Number of stemmed documents: 854

First 10 stemmed documents:

Document 8334487:
['biomimet', 'built', 'integr', 'specif', 'membran', 'artifici', 'synthet', 'r

Document 8334505:
['zoledron', 'acid', 'zol', 'adjuv', 'breast', 'suggest', 'zol', 'associ', 'in

Document 8334684:
['tripl', 'neg', 'breast', 'tnbc', 'deadliest', 'form', 'breast', 'success',

Document 8334718:
['lung', 'remain', 'lead', 'relat', 'death', 'unit', 'state', 'although', 'alv

Document 8334819:
['properti', 'platinum', 'render', 'desir', 'diagnosi', 'detect', 'surgeri',

Document 8335232:
['select', 'plasmon', 'critic', 'penetr', 'clinic', 'applic', 'photoacoust',

Document 8335457:
['patient', 'malign', 'ascit', 'ma', 'display', 'sever', 'symptom', 'pain', 'a

Document 8335543:
['iron', 'oxid', 'found', 'widespread', 'applic', 'differ', 'area', 'includ',

Document 8335571:
['previou', 'identifi', 'relev', 'gene', 'signal', 'pathway', 'hamper', 'human

Document 8336594:
['kind', 'ray', 'comput', 'tomographi', 'ct', 'fluoresc', 'dual', 'modal', 'na
```

```
cleaned_df = pd.DataFrame(list(zip(article_id, text)), columns = ['Article ID', 'Cl
```

```
cleaned_df.to_csv('/content/grive/MyDrive/ML Literature Review/Chloe Brown/Revisiti
```

### Step 3 - Construct a Document-Term Matrix

A document-term matrix determines how often each word occurs in each document.

We will be using the corpora module from the gensim package. The models module wil also be imported and will be used later on. Documentation/API Reference: https://radimrehurek.com/gensim/apiref.html

Steps:

1. Create the Dictionary
2. Convert Dictionary to Bag-of-Words

##### Step 3.1 - Create the Dictionary

In this step, the dictionary will be created. The dictionary contains the unique words in the data set. To do this, the `corpora.Dictionary` class will be used. Documentation: https://radimrehurek.com/gensim/corpora/dictionary.html

The `corpora.Dictionary` constructor creates a mapping between words and their integer ids. This will be assigned to the variable `dictionary`. It takes in the paramter `documents`, which must be a list of lists of strings, as is the case in the cleaned dataset above.

The other parameter, `prune_at`, which sets a limit on how many words the dictionary will keep, will not be used.

The variable `vocabulary` will represent the number of unique words in dataset.

`dictionary` has the attribute `token2id` which shows the mapping of words to unique integer ids.

```
from gensim import corpora, models

# Create the dictionary
dictionary = corpora.Dictionary(text)

# Define the vocabulary (= number unique words in dataset)
vocabulary = len(dictionary)

print("Unique words in dataset: %d words" %vocabulary)
```

```
Unique words in dataset: 6999 words
```

##### Step 3.2 - Convert Dictionary to Bag-of-Words

To convert the dictionary to a bag-of-words, the function `doc2bow()` will be used. (For link to documentation see Step 3.1).

This function returns a list of vectors. The length of the list is equal to the number of documents in the dataset, which each vector representing one document. Each vecor is a list of tuples. The first term in the tuple is an integer id corresponding to a word according to the mapping in `dictionary`. The second term in the tuple is the frequency with which the word associated with that id occurs in the document. That is, each tuple contains the following information:

(word ID, word frequency in document n)

Note that words that do not appear in a given document will not appear in that document's vector.

The `document` parameter for `doc2bow` is used in this project. `document` is a list of strings. This will be an individual document from the cleaned dataset from Step 2. Thus, the cleaned dataset must be looped through so this can be applied to each document.

The bag-of-words will be assigned to the variable `corpus`.

The paramter `allow_update` will be left at its default value (false). This parameter would allow `dictionary` to update itself if it came across new tokens if it were set to true.

The paramter `return_missing` will also be left at its default value (false). If it were set to true, it would return a list of tokens prersent in the document but not in the dictionary.

```
# Define list for corpus
corpus = []

# Loop through texts
# doc represents one document
for doc in text:

  # Create bag of words for doc and add it to the corpus
  corpus.append(dictionary.doc2bow(doc))

print("First 10 documents as Bag-of-Words:")
for i in range(10):
  print("\nDocument %d:" %article_id[i])
  print(corpus[i])
```

```
First 10 documents as Bag-of-Words:

Document 8334487:
[(0, 1), (1, 1), (2, 1), (3, 1), (4, 1), (5, 1), (6, 1), (7, 1), (8, 1), (9,

Document 8334505:
[(81, 1), (82, 1), (83, 1), (84, 1), (85, 3), (86, 1), (87, 1), (88, 1), (89,

Document 8334684:
[(20, 1), (40, 1), (46, 1), (57, 1), (69, 1), (84, 2), (89, 1), (91, 2), (118

Document 8334718:
[(24, 3), (33, 1), (41, 1), (46, 2), (61, 1), (72, 1), (80, 2), (99, 1), (100

Document 8334819:
[(21, 2), (35, 1), (40, 1), (46, 1), (48, 1), (49, 1), (53, 4), (62, 2), (69,

Document 8335232:
[(4, 1), (12, 1), (20, 1), (24, 1), (25, 1), (30, 1), (46, 1), (57, 1), (77,

Document 8335457:
[(35, 1), (46, 2), (60, 1), (85, 1), (89, 1), (99, 1), (104, 1), (108, 1), (1

Document 8335543:
[(1, 1), (15, 1), (16, 6), (40, 1), (42, 2), (44, 1), (52, 1), (53, 1), (62,

Document 8335571:
[(19, 1), (24, 2), (39, 1), (45, 1), (46, 2), (54, 1), (57, 1), (59, 1), (67,

Document 8336594:
[(21, 1), (90, 1), (91, 1), (117, 1), (125, 1), (135, 1), (140, 2), (153, 5),
```

#### Step 5 - Create the Model

We will use the Gensim ensembleLDA algorithm. The model will then be converted to a standard LDA model for the purposes of analysis.

- ensembleLDA: https://radimrehurek.com/gensim/models/ensemblelda.html
- Standard LDA: https://radimrehurek.com/gensim/models/ldamodel.html

```
from gensim.models import EnsembleLda

alpha = 0.1
beta = 0.1

elda = elda = EnsembleLda(topic_model_class = 'lda',
                          corpus=corpus,
                          id2word=dictionary,
                          num_models=50,
                          alpha = alpha,
                          eta = beta,
                          passes = 20)
```

```
WARNING:gensim.models.ldamodel:too few updates, training might not converge; 
```

```
elda.recluster(eps = 0.2)
lda_final = elda.generate_gensim_representation()
lda_final.print_topics()
```

```
[(0,
  '0.001*"nh" + 0.001*"kill" + 0.000*"biocarri" + 0.000*"stem" + 0.000*"make"
+ 0.000*"royal" + 0.000*"3o" + 0.000*"chemistri" + 0.000*"vulner" +
0.000*"click"'),
 (1,
  '0.038*"tam" + 0.023*"m2" + 0.017*"m1" + 0.017*"associ" + 0.015*"phenotyp"
+ 0.015*"polar" + 0.013*"microenviron" + 0.012*"anti" + 0.012*"induc" +
0.011*"growth"'),
 (2,
  '0.073*"exosom" + 0.048*"mir" + 0.020*"deriv" + 0.013*"eoc" +
0.011*"express" + 0.011*"m2" + 0.009*"polar" + 0.009*"hypox" + 0.009*"mirna"
+ 0.009*"microenviron"'),
 (3,
  '0.056*"micel" + 0.048*"dox" + 0.014*"releas" + 0.014*"mcf" + 0.010*"free"
+ 0.009*"ml" + 0.009*"copolym" + 0.009*"block" + 0.009*"mpeg" +
0.009*"adr"'),
 (4,
  '0.089*"cpg" + 0.038*"odn" + 0.009*"enhanc" + 0.009*"activ" +
0.009*"oligodeoxynucleotid" + 0.008*"immunotherapi" + 0.007*"liposom" +
0.007*"induc" + 0.007*"respons" + 0.007*"contain"'),
 (5,
  '0.038*"imag" + 0.018*"magnet" + 0.018*"iron" + 0.014*"mri" + 0.013*"oxid"
+ 0.011*"contrast" + 0.011*"spion" + 0.010*"agent" + 0.010*"mr" +
0.009*"enhanc"')]
```

```
lda_final.save('')
```

```
from gensim.models import CoherenceModel

# create the coherence model
coherence_model_lda = CoherenceModel(model = lda_final, texts = text, dictionary=di

# get the coherence score
coherence_score = coherence_model_lda.get_coherence()

print('Final coherence score (Cv): %f' %coherence_score)
```

```
Final coherence score (Cv): 0.589477
```

```
coherence_model_lda.get_coherence_per_topic()
```

```
[0.4130806884722843,
 0.624529044710907,
 0.6803449318603165,
 0.7533153597551968,
 0.4803173307484706,
 0.585277421929308]
```

### Step 6 - Create the Figures

#### Figure 3 - pyLDAvis

```
import pyLDAvis.gensim_models
pyLDAvis.enable_notebook()
vis = pyLDAvis.gensim_models.prepare(lda_final, corpus, dictionary=lda_final.id2wor
vis
```

```
/usr/local/lib/python3.7/dist-packages/pyLDAvis/_prepare.py:247: FutureWarning
  by='saliency', ascending=False).head(R).drop('saliency', 1)
```

Selected Topic: 0 Previous Topic Next Topic Clear Topic

Intertopic Distance Map (via multidimensional scaling)

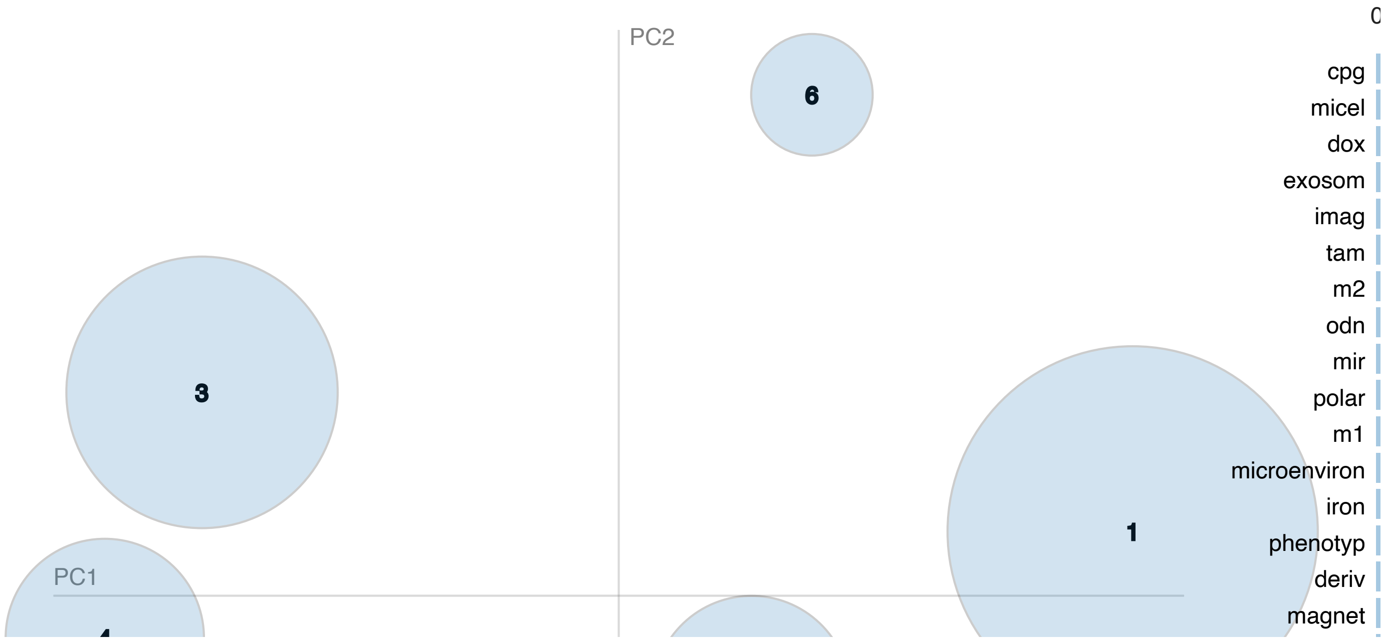

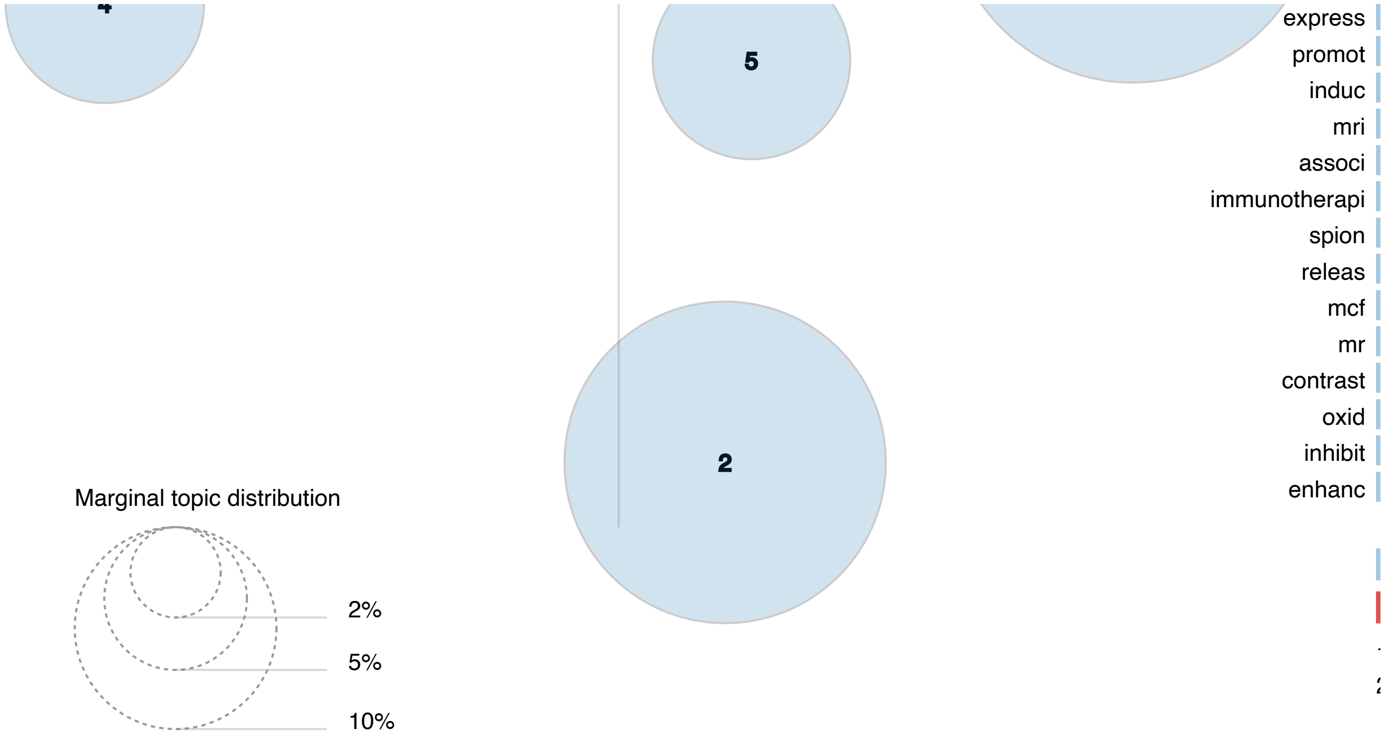


#### Figure 4 - Word Clouds and Heat Maps

```
import matplotlib.pyplot as plt
from matplotlib import colormaps, rcParams
import pandas as pd
from wordcloud import WordCloud, get_single_color_func


# Citation: https://amueller.github.io/word_cloud/auto_examples/colored_by_group.ht
class SimpleGroupedColorFunc(object):
    """Create a color function object which assigns EXACT colors
       to certain words based on the color to words mapping

       Parameters
       ----------
       color_to_words : dict(str -> list(str))
         A dictionary that maps a color to the list of words.

       default_color : str
         Color that will be assigned to a word that's not a member
         of any value from color_to_words.
    """

    def __init__(self, color_to_words, default_color):
        self.word_to_color = {word: color
                              for (color, words) in color_to_words.items()
                              for word in words}

        self.default_color = default_color

    def __call__(self, word, **kwargs):
        return self.word_to_color.get(word, self.default_color)
```

```
# get word frequency dictionary for each topic
freq_df = pd.read_csv("Living_review/living_review_word_cloud.csv")

# topic 0
keys0 = list(freq_df['Top Words (Topic 0)'].dropna())
vals0 = list(freq_df['Word Frequency (Topic 0)'].dropna())
words0 = dict(zip(keys0, vals0))

# topic 1
keys1 = list(freq_df['Top Words (Topic 1)'].dropna())
vals1 = list(freq_df['Word Frequency (Topic 1)'].dropna())
words1 = dict(zip(keys1, vals1))

# topic 2
keys2 = list(freq_df['Top Words (Topic 2)'].dropna())
vals2 = list(freq_df['Word Frequency (Topic 2)'].dropna())
words2 = dict(zip(keys2, vals2))

# topic 3
keys3 = list(freq_df['Top Words (Topic 3)'].dropna())
vals3 = list(freq_df['Word Frequency (Topic 3)'].dropna())
words3 = dict(zip(keys3, vals3))

# topic 0
keys4 = list(freq_df['Top Words (Topic 4)'].dropna())
vals4 = list(freq_df['Word Frequency (Topic 4)'].dropna())
words4 = dict(zip(keys4, vals4))

# topic 5
keys5 = list(freq_df['Top Words (Topic 5)'].dropna())
vals5 = list(freq_df['Word Frequency (Topic 5)'].dropna())
words5 = dict(zip(keys5, vals5))# topic 0 wordcloud (with colors)
```

```
# define dictionary for colors
color_df = pd.read_csv("Living_review/living_colors.csv")
blue = list(color_df['blue'].dropna())
green = list(color_df['green'].dropna())
orange = list(color_df['orange'].dropna())
red = list(color_df['red'].dropna())
purple = list(color_df['purple'].dropna())
gray = list(color_df['gray'].dropna())
color_dict = {(86, 180, 233):blue, (213, 94, 0):green, (0, 0, 0):orange, (100, 117,
default_color = (230, 159, 0)

# Create a color function with multiple tones
grouped_color_func = SimpleGroupedColorFunc(color_dict, default_color)
```

Example Word Cloud (From Living Review Topic 1 Figure)

```
# topic 1 wordcloud (with colors)
wc1 = WordCloud(background_color = "white").generate_from_frequencies(words1)
wc1.recolor(color_func = grouped_color_func)
plt.imshow(wc1)
plt.axis("off")
plt.show()
```

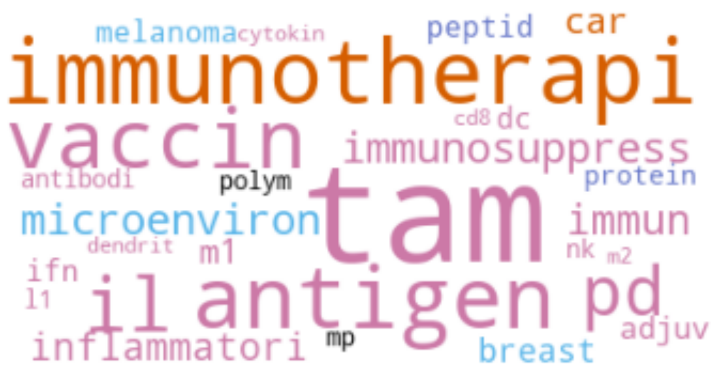


#### Step 7 - Living Review

To create a "living review", topics were extracted from an additional set of documents.

```
from gensim.corpora.dictionary import Dictionary
from gensim.models.ldamodel import LdaModel
```

```
import pandas as pd

# Define file path
path1 = ''
path2 = ''

# Import data frame
df1 = pd.read_csv(path1)
df2 = pd.read_csv(path2)



# get unseen documents
df_help = df1.merge(df2, how = "outer", on = "Article ID", indicator = True)
df = df_help[df_help["_merge"] == 'left_only']

# should be 77 rows long
df.reset_index(drop = True)
```

| | Article ID | Article URL_x | Title_x | Journal_x | [illegible] |
|---|---|---|---|---|---|
| **0** | 90543108 | https://sysrev.com/p/43536/article/90543108 | Hierarchically targetable fiber rods decorated... | Acta Biomaterialia | [illegible] |
| **1** | 90543116 | https://sysrev.com/p/43536/article/90543116 | CD73 specific siRNA loaded chitosan lactate na... | J Control Release | |
| **2** | 90543117 | https://sysrev.com/p/43536/article/90543117 | In situ vaccination with cowpea mosaic virus n... | Nature Nanotechnology | [illegible] |
| **3** | 90543118 | https://sysrev.com/p/43536/article/90543118 | Core–Shell Distinct Nanodrug Showing On-Demand... | ACS Nano | [illegible] |
| | | | Cell Membrane [illegible] | | |

| | | | | |
|---|---|---|---|---|
| **4** | 90543119 | https://sysrev.com/p/43536/article/90543119 | Cell-Membrane Immunotherapy Based on Natural K... | ACS Nano |
| **...** | ... | ... | ... | ... |
| **72** | 90543366 | https://sysrev.com/p/43536/article/90543366 | ADJUVANT PROPERTIES OF NANOPARTICLES IMMOBILIZ... | Biotechnologia Acta |
| **73** | 90543369 | https://sysrev.com/p/43536/article/90543369 | Co-delivery of RNAi and chemokine by polyargin... | J Control Release |
| **74** | 90543374 | https://sysrev.com/p/43536/article/90543374 | PET Imaging of Tumor-Associated Macrophages wi... | Journal of Nuclear Medicine |
| **75** | 90543375 | https://sysrev.com/p/43536/article/90543375 | Delivery of nitric oxide with a | Nature |

```
# https://www.geeksforgeeks.org/pandas-get-the-elements-of-series-that-are-not-pres
df = df1[~df1['Article ID'].isin(df2['Article ID'])]
df
```

| | Article ID | Article URL | Title | Journal |
|---|---|---|---|---|
| **854** | 90543108 | https://sysrev.com/p/43536/article/90543108 | Hierarchically targetable fiber rods decorated... | Acta Biomaterialia |
| **855** | 90543116 | https://sysrev.com/p/43536/article/90543116 | CD73 specific siRNA loaded chitosan lactate na... | J Control Release |

| | | | | |
|---|---|---|---|---|
| **856** | 90543117 | https://sysrev.com/p/43536/article/90543117 | In situ vaccination with cowpea mosaic virus n... | Nature Nanotechnology |
| **857** | 90543118 | https://sysrev.com/p/43536/article/90543118 | Core–Shell Distinct Nanodrug Showing On-Demand... | ACS Nano |
| **858** | 90543119 | https://sysrev.com/p/43536/article/90543119 | Cell-Membrane Immunotherapy Based on Natural K... | ACS Nano |
| **...** | ... | ... | ... | ... |
| **926** | 90543366 | https://sysrev.com/p/43536/article/90543366 | ADJUVANT PROPERTIES OF NANOPARTICLES IMMOBILIZ... | Biotechnologia Acta |
| **927** | 90543369 | https://sysrev.com/p/43536/article/90543369 | Co-delivery of RNAi and chemokine by polyargin... | J Control Release |
| **928** | 90543374 | https://sysrev.com/p/43536/article/90543374 | PET Imaging of Tumor-Associated Macrophages wi... | Journal of Nuclear Medicine |
| **929** | 90543375 | https://sysrev.com/p/43536/article/90543375 | Delivery of nitric oxide with a nanocarrier pr... | Nature Nanotechnology |

```
# Get abstracts
document_set = df['Abstract']
print('Abstracts:')
document_set = document_set.reset_index(drop = True)
print(document_set)

# Get article ID's
article_id = df['Article ID']
print('\nArticle ID:')
article_id = article_id.reset_index(drop = True)
print(article_id)

# Compile features of dcument set
N_docs = len(document_set)

print("\nNumber of documents: %d documents" %N_docs)
```

```
Abstracts:
0     The shapes of drug carriers have significant e...
1     The efficacy of conventional anti-tumor immuno...
2     Nanotechnology has tremendous potential to con...
3     Among various inflammatory factors/mediators, ...
4     Developing effective immunotherapies with low ...
                              ...
72    The aim of the research was to compare the cha...
73    Myeloid-Derived Suppressor Cells (MDSCs), immu...
74    Tumor-associated macrophages (TAMs) are increa...
75    Abnormal tumour vasculature has a significant ...
76    Although radiotherapy has been established as ...
Name: Abstract, Length: 77, dtype: object

Article ID:
0     90543108
1     90543116
2     90543117
3     90543118
4     90543119
        ...
72    90543366
73    90543369
74    90543374
75    90543375
76    90543377
Name: Article ID, Length: 77, dtype: int64

Number of documents: 77 documents
```

Documents were cleaned using the code from Step 2.

```
# convert documents to bag of words model and get topics
bow = []
topics = []
for i in range(len(cleaned_df)):
  curr_bow = lda.id2word.doc2bow(cleaned_df['Cleaned Text'][i])
  bow.append(curr_bow)
  topics.append(lda.get_document_topics(curr_bow))

cleaned_df['BOW'] = bow
cleaned_df['Topics'] = topics
```

Colab paid products - Cancel contracts here
